# Simulations provide a mechanistic interpretation of Bateman gradients

**DOI:** 10.1101/2024.12.03.626594

**Authors:** Krish Sanghvi, Jonathan M. Henshaw, Alex Kacelnik, Tim Janicke, Irem Sepil

**Affiliations:** Department of Biology, University of Oxford, U.K.; Institute of Biology, University of Freiburg, Germany; CNRS Centre for Functional and Evolutionary Ecology, Montpellier, France

**Keywords:** cryptic female choice, polygyny, polyandry, selection gradient, sexual selection, sperm competition

## Abstract

The Bateman gradient (BG or 𝛽_𝑠𝑠_) is a fundamental metric of sexual selection, and is often interpreted as the fitness advantage individuals can gain by increasing their number of mates. However, researchers commonly misconceive the relationship between mating success (MS) and reproductive success (RS) as always being causal, leading to a misinterpretation of the BG. The BG is phenomenological and therefore uninformative about the mechanisms leading to correlations between MS and RS. Here, we simulate sequential matings in males from biological systems with varying levels of anisogamy and sperm competition, to better understand the underlying processes that modulate male BGs. We show that the BG does not solely reflect the causal influence of MS on RS as is typically interpreted, but is also a consequence of covariances between male MS and male ejaculate traits, female fecundity, or female post-mating decisions. For example, we show that the BG is substantially shallower when the ejaculate size of males negatively co-varies with their MS, such as in species with alternative male reproductive tactics. Such covariances can lead to a misinterpretation of the measured BG, because here, it does not merely represent the strength of selection on intra-sexual competition for mates. We suggest how such covariances can be visually assessed, and accounted for using partial Bateman gradients, to better interpret the influence of MS on RS, thus the strength of pre-copulatory sexual selection. Among other simplifications, our simulations have a male-only focus, but they provide insights into the mechanisms driving inter- and intra-specific variation in the BG, clarify its meaning, and improve our understand of sexual selection.

## Introduction

The Bateman gradient (Bateman, 1948; henceforth BG or 𝛽_𝑠𝑠_) is a cornerstone of sexual selection theory (Arnold and Duvall, 1994; Henshaw and Jones, 2019). Typically, the BG is measured as the slope of the ordinary least-square (OLS) linear regression of the relativised number of offspring produced (i.e. reproductive success: RS) on the relativised number of mates (i.e. mating success: MS) (Anthes et al, 2017; Arnold, 1994; Jones, 2009; Wade and Shuster, 2005). Multiple studies on wild (e.g. Mobley and Jones, 2007; Rios-Cardenas, 2005; Rodriguez-Munoz et al, 2010; Schulte-Hostedde et al, 2004; Webster et al, 2007) and lab animals (e.g. Andrade, 2005; Bussiere et al, 2008; Davies et al, 2023; Fritzsche et al, 2013; Jones et al, 2002, 2004; Pelissie et al, 2012), as well as plants (Tonnabel et al, 2019) have demonstrated the usefulness of the BG in understanding pre-copulatory sexual selection. Specifically, the BG is used to interpret the selective fitness advantage gained from intra-sexual competition for mates. Due to males being less gamete limited than females, male BGs are typically positive, and steeper than that of females (Janicke et al, 2016). Furthermore, males have greater variance in their RS and MS compared to females. These observations have led to the predictions of stronger strength of and opportunity for pre-copulatory sexual selection on males than females. However, not all studies agree with these predictions (e.g. Apakupakul et al, 2015; Douglas et al, 2020; Gopurenko et al, 2007; Jones et al, 2000; Nason and Kelly, 2020). Additionally, recent studies argue that the BG is often misinterpreted (Anthes et al, 2017; Collet et al, 2014; Klug et al, 2010; Tang-Martinez, 2016), because of its failure to partition pre-from post-copulatory episodes of sexual selection, or reveal what specific processes generate correlations between MS and RS (Arnold, 1994; Collet et al, 2014; Fromonteil et al, 2023; Tang-Martinez, 2006; 2016).

Within a sex, BG measures the statistical relationship between an individual’s MS and RS. This relationship is interpreted as the potential fitness benefits gained by individuals, when they acquire more mates. However, there are very few empirical tests that experimentally manipulate MS to test its causal influence on RS (but see Andrade and Kasumovic, 2005). Instead, most studies, including Bateman’s (1948), use standing variation in MS and RS to assess their relationship (e.g. Mobley and Jones, 2007; Rios-Cardenas, 2005; Rodriguez-Munoz et al, 2010; Schulte-Hostedde et al, 2004; Webster et al, 2007). Here, however, the correlation between MS and RS is not just the consequence of a causal effect of MS on RS while holding other variables constant, but can partly arise due to MS co-varying with other traits (e.g. female traits) that influence RS (Anthes et al, 2010; Davies et al, 2023; Evans and Garcia-Gonzalez, 2016; Henshaw et al, 2018; McDonald and Pizzari, 2016; Parker and Tang-Martinez, 2005; Figure 1). For instance, some studies show that dominant males mate with more fecund (Collet et al, 2014; Gerlach et al, 2012; Kraak and Bakker, 1998) or less promiscuous females (Greenway et al, 2021). This can lead to dominant males, who are usually better at mate acquisition than subordinate males, having a higher RS partly due to mating with higher quality females or less promiscuous females, rather than just due to mating with more females. Likewise, females might favour sperm of dominant or ornamented males compared to subordinate or less ornamented males (Dean et al, 2011; Pizzari and Birkhead, 2000). Here, similarly, dominant/ornamented males would gain higher RS not only due to mating with more females, but also due to a higher paternity share with each female. Similarly, females allocating a larger proportion of their available eggs to be fertilised by more dominant males (Horvathova et al, 2012; Skinner and Watt, 2007), might further lead to a confounded positive relationship between MS and RS. Unless the influence of such male-female interactions on the BG are explicitly understood, we cannot interpret the measured BG as merely representing the strength of pre-copulatory sexual selection for mates.

**Figure 1:**
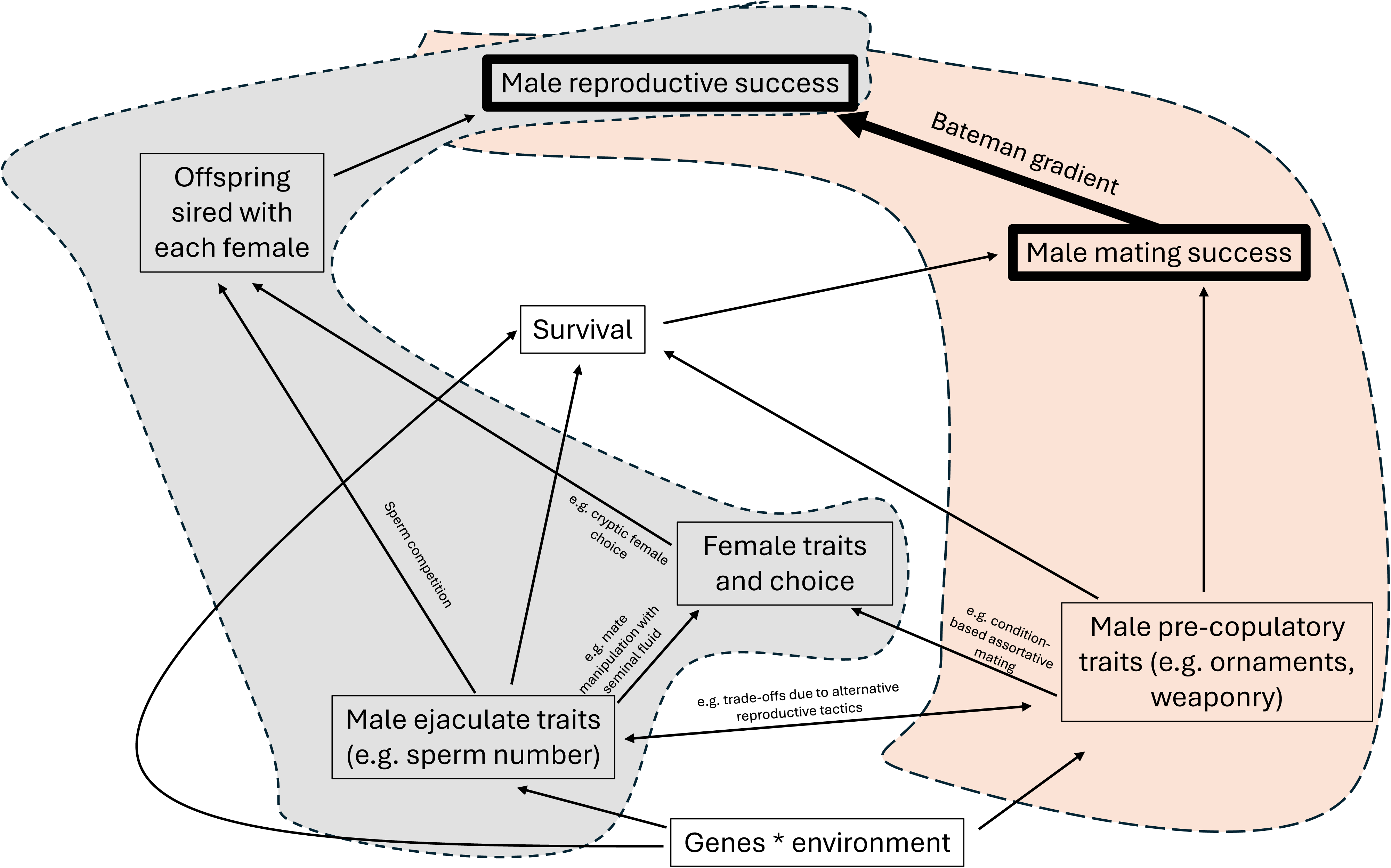
Example of how male mating success (MS), other traits, and reproductive success (RS) might interact. The BG is often interpreted as the causal influence of varying MS on RS, reflecting the strength of pre- copulatory sexual selection acting on competition for mates. However, the relationship between MS on RS need not always be causal. MS can correlate with RS due to covariances between MS and other reproductive traits of both males and females. These covariances might inflate or deflate our interpretation of the strength of sexual selection. When such confounding occurs, interpretations of the resultant BG as the strength of sexual selection on mates can be misleading. Boxes inside the orange highlighted areas are typically interpreted as pre- copulatory sexual selection; those inside the grey areas are interpreted as post-copulatory sexual selection. Arrows show likely direction of causality, and can represent positive or negative effects. Figure adapted from Anthes et al (2010). Similar path diagrams can be constructed for females, whereby female MS correlates with RS through its covariances with other male or female traits.

The BG does not partition selection on post-copulatory traits, from selection on pre-copulatory traits that improve MS (Collet et al, 2012; Davies et al, 2023; Devigili et al, 2015; Evans and Garcia-Gonzalez, 2016; Krakauer, 2008; Pelissie et al, 2014). In males, MS is typically a consequence of the expression of traits such as male social status, weaponry, lifespan, mate-finding ability and latency, attractiveness, mate-coercion ability, and male or female choosiness (Arnold, 1995; Collet et al, 2014; Douglas et al, 2020; Gao et al, 2020; Gowaty and Hubbell, 2005; Tang-Martinez, 2005; 2016). The numbers of offspring a male produces with each female (the cumulative of which is RS) or his paternity share, are also a consequence of the expression of other reproductive traits (Anthes et al, 2017; Pischedda and Rice, 2012). These could include male traits such as ejaculate quality and quantity, sperm competitiveness, ejaculate allocation strategies, mate guarding, or paternal investment, as well as female traits such as egg number, egg allocation, degree of cryptic female choice, polyandry, or maternal investment. Distinguishing the influence of male ejaculate and female post-mating traits on reproductive RS, independent of MS, is crucial for determining whether the BG reflects only precopulatory sexual selection on mate acquisition or also postcopulatory sexual selection (Davies et al, 2023; Evans and Garcia-Gonzalez, 2016; Marie-Orleach et al, 2024). For instance, male mating success might co-vary negatively with male ejaculate size (Gowaty and Hubbell, 2005), such as in species with alternative male reproductive tactics whereby dominant males, who are better at monopolizing females (and thus have higher MS), allocate small ejaculates to each female. In the presence of sperm competition, such differential ejaculate allocation might lead to a lower paternity share for dominant compared to subordinate males, who mate with fewer females but allocate larger ejaculates to each female (e.g. Alonzo and Warner, 2000; Durant et al, 2016; Froman et al, 2002; Gage et al, 1995; Klaus et al, 2011; Lupold et al, 2014; Preston et al, 2001; Saunders and Shuster, 2020; Warner et al, 1995). Such negative covariances (e.g. Kvanermo and Simmons, 2013; Parker et al, 2013) might lead to underestimating the fitness benefits gained by increasing the number of mating partners. Such covariances might explain why subordinate males in species with alternative reproductive tactics often have similar lifetime reproductive success compared to dominant males, despite having lower mating success (reviewed in Da Silva et al, 2024). Male pre- and post-copulatory traits could also co-vary positively due to condition dependence (e.g. Bath et al, 2023; De Nardo et al, 2021; Devigili et al, 2013; Janicke and Chapuis, 2016; reviewed in Simmons et al, 2017). For instance, individuals in higher condition might develop better pre-and post-copulatory traits than individuals in lower condition (Hare and Simmons, 2022; Janicke et al, 2015; Mehlis et al, 2015; Morimoto et al, 2016). Under such covariances, the BG would represent the strength of both, pre- and post-copulatory sexual selection, rather than just pre-copulatory selection for more mates (i.e. its typical interpretation).

Recording the specific mating order in which individuals encounter mates in a mating sequence might allow us to assess whether covariances between male MS and other traits exist (e.g. Pelissie et al, 2014). For instance, Sanghvi et al (2024a) find in *D. melanogaster* that males with higher MS produce more offspring with females early in the mating sequence, but produce fewer offspring with females later in their mating sequence, compared to males with lower MS. This pattern, of males with higher MS having higher intercepts but steeper slopes for offspring production with each female through the mating sequence, might be caused by male MS covarying positively with male ejaculate allocation. If the exact mating order of females is known, assessing the patterns of offspring production through the mating sequence might allow inferring the type of covariances between MS and other variables, and the magnitude of their effect on the BG. However, inferring the presence and magnitude of covariances by recording the exact mating order has not been demonstrated before.

We use simulations to investigate the potential mechanisms driving variation in the male Bateman gradient (BG), and how these might distort the interpretation of the strength of pre-copulatory sexual selection. Specifically, we aim to disentangle the relationship between male mating success (MS) and reproductive success (RS), by exploring how covariances between male MS and other male or female reproductive traits and post-mating decisions contribute to variation in the BG. Importantly, we simulate mating sequences to assess whether tracking the order in which females encounter males improves our understanding of the BG. Previous studies have reviewed and provided verbal models or experimental tests for how some covariances might influence the BG (e.g. Anthes et al, 2010, 2017; Evans and Garcia-Gonzalez, 2016; Henshaw et al, 2018; McDonald and Pizzari, 2015). Here, we include a broader range of covariance scenarios and place emphasis on the usefulness of recording the mating sequence. It is often assumed that the strength of pre-copulatory sexual selection for mate acquisition is stronger in high than low anisogamy species, for the gamete- abundant sex (Henshaw et al, 2022; Janicke, 2024; Jones et al., 2002; Lehtonen, 2022; Lehtonen & Parker, 2024). However, whether sperm competition can modulate the influence of anisogamy on the BG remains unclear. Therefore, we consider biological systems with varying degrees of anisogamy and sperm competition to provide a more generalizable understanding of the BG across taxa. While our simulations focus on males, the principles highlighted herein are applicable to females too.

## Methods

We used agent-based simulations to investigate how covariances between male MS and various other male or female reproductive traits affect RS, thus modulating the BG. We modelled the number of offspring produced by individual focal males interacting sequentially with multiple females, and subsequently calculated male BGs. All analyses were conducted in R.v.4.3.2 (R core team, 2020). Simulation code and simulated data can be found at Open science framework DOI:10.17605/OSF.IO/WPE8M.

### Simulation overview

We simulated nine covariance scenarios across four different biological systems corresponding to each combination of high or low anisogamy, with or without sperm competition. Detailed descriptions of all scenarios are provided below. In all simulations, male reproductive success was modelled to depend on the following traits (also see Table 1):

**Table 1:**
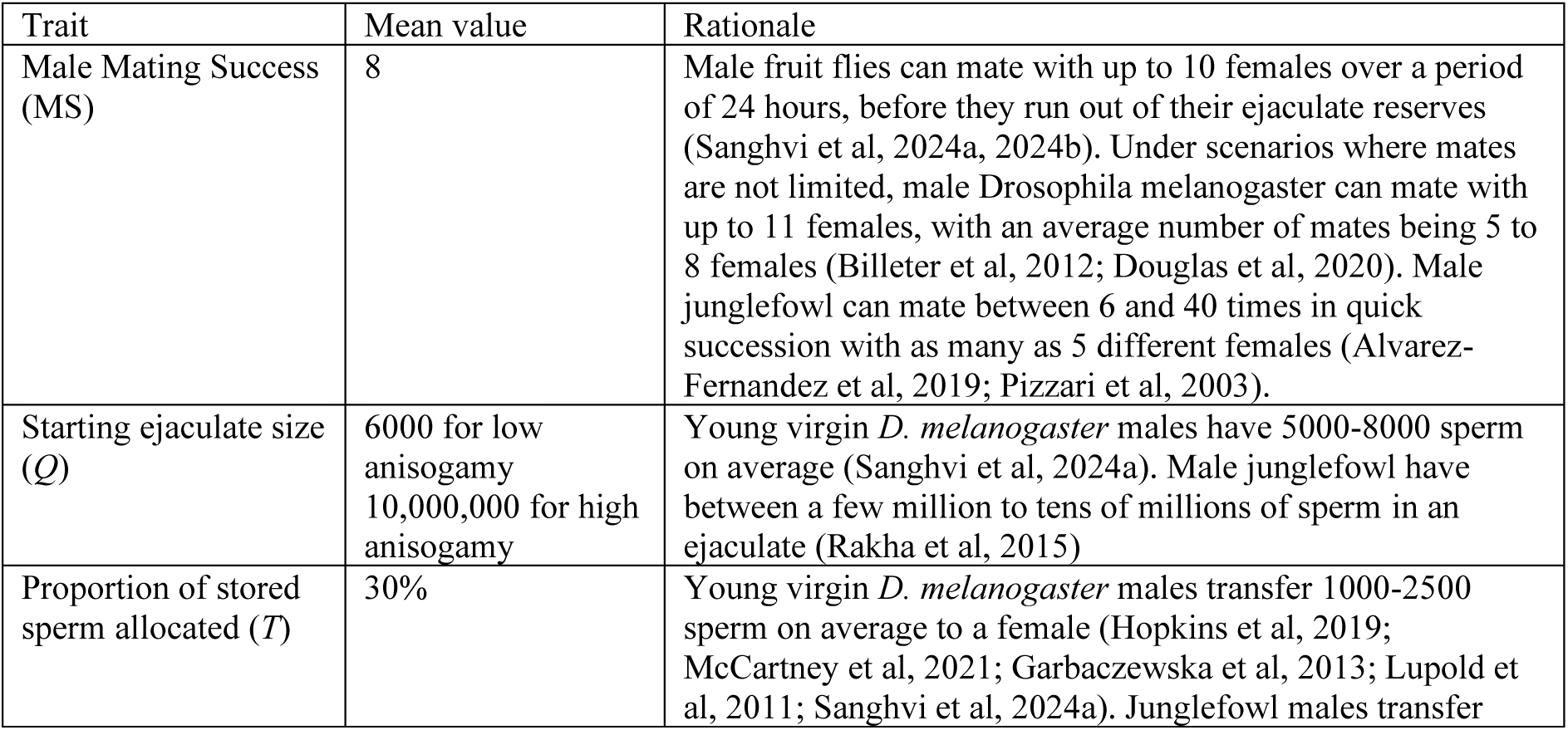
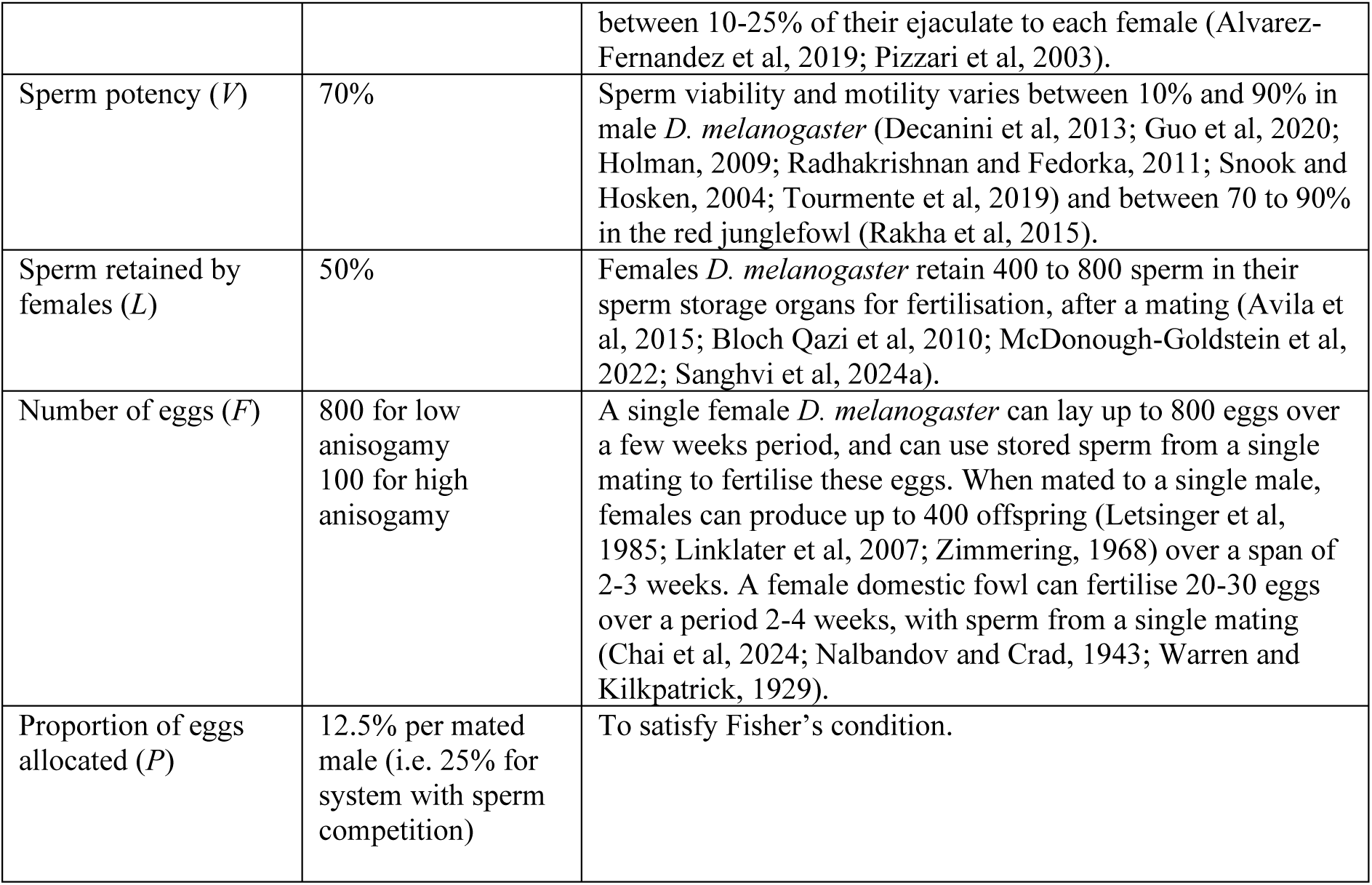
mean values for creating normal distributions used for sampling each trait in our simulations for focal males and females, and the rationale for using these. To allow comparison between the low and high anisogamy systems, we retained the same trait values and distributions for all variables in both anisogamy systems, except for starting ejaculate size (*Q*) and number of eggs (*F*).

| Trait | Mean value | Rationale |
| --- | --- | --- |
| Male Mating Success (MS) | 8 | Male fruit flies can mate with up to 10 females over a period of 24 hours, before they run out of their ejaculate reserves (Sanghvi et al, 2024a, 2024b). Under scenarios where mates are not limited, male *Drosophila melanogaster* can mate with up to 11 females, with an average number of mates being 5 to 8 females (Billeter et al, 2012; Douglas et al, 2020). Male junglefowl can mate between 6 and 40 times in quick succession with as many as 5 different females (Alvarez-Fernandez et al, 2019; Pizzari et al, 2003). |
| Starting ejaculate size ($Q$) | 6000 for low anisogamy<br>10,000,000 for high anisogamy | Young virgin *D. melanogaster* males have 5000-8000 sperm on average (Sanghvi et al, 2024a). Male junglefowl have between a few million to tens of millions of sperm in an ejaculate (Rakha et al, 2015) |
| Proportion of stored sperm allocated ($T$) | 30% | Young virgin *D. melanogaster* males transfer 1000-2500 sperm on average to a female (Hopkins et al, 2019; McCartney et al, 2021; Garbaczewska et al, 2013; Lupold et al, 2011; Sanghvi et al, 2024a). Junglefowl males transfer |
|  |  | between 10-25% of their ejaculate to each female (Alvarez-Fernandez et al, 2019; Pizzari et al, 2003). |
| Sperm potency ($V$) | 70% | Sperm viability and motility varies between 10% and 90% in male *D. melanogaster* (Decanini et al, 2013; Guo et al, 2020; Holman, 2009; Radhakrishnan and Fedorka, 2011; Snook and Hosken, 2004; Tourmente et al, 2019) and between 70 to 90% in the red junglefowl (Rakha et al, 2015). |
| Sperm retained by females ($L$) | 50% | Females *D. melanogaster* retain 400 to 800 sperm in their sperm storage organs for fertilisation, after a mating (Avila et al, 2015; Bloch Qazi et al, 2010; McDonough-Goldstein et al, 2022; Sanghvi et al, 2024a). |
| Number of eggs ($F$) | 800 for low anisogamy<br>100 for high anisogamy | A single female *D. melanogaster* can lay up to 800 eggs over a few weeks period, and can use stored sperm from a single mating to fertilise these eggs. When mated to a single male, females can produce up to 400 offspring (Letsinger et al, 1985; Linklater et al, 2007; Zimmering, 1968) over a span of 2-3 weeks. A female domestic fowl can fertilise 20-30 eggs over a period 2-4 weeks, with sperm from a single mating (Chai et al, 2024; Nalbandov and Crad, 1943; Warren and Kilpatrick, 1929). |
| Proportion of eggs allocated ($P$) | 12.5% per mated male (i.e. 25% for system with sperm competition) | To satisfy Fisher's condition. |

#### Male traits

- Male mating success (MS): the total number of females a male mated with. MS is often determined by the expression of male weaponry, male ornamentation, mate coercion ability, energetic and physiological mating costs, courtship ability, or female choosiness. However, we did not simulate these underlying traits explicitly.
- Starting ejaculate size (*Q*): the number of sperm a male has available in his ejaculate reserves before encountering any females. The starting ejaculate size could also represent starting seminal fluid quantity.
- Sperm potency (*V*): the proportion of sperm in the male’s ejaculate that has the capacity to fertilize an egg and produce offspring due to intrinsic sperm quality of the donor. Biologically, *V* might represent sperm viability, egg penetration ability, or sperm DNA integrity. *1-V* is the proportion of a male’s sperm incapable of fertilisation or offspring production.
- Ejaculate allocation (*T*): the proportion of a male’s remaining sperm that he transfers to a given female in his mating sequence when copulating.

#### Female traits

- Number of eggs (F): the number of eggs a female has available that are viable and can produce offspring, if fertilised.
- Proportion of eggs allocated (P): the proportion of her available eggs the female allocates to be fertilised by the current male she is copulating with.
- Sperm retained (L): the proportion of a male’s inseminated sperm that a female uses (i.e. does not eject or absorb, and retains for fertilisation). In some cases, this trait could also be a function of a sperm’s ability to enter female sperm storage organs.

### Simulated scenarios

We simulated nine different scenarios for each of the four biological systems. In the null scenario, the relationship between MS and RS was causal, and male mating success did not co-vary with any other male or female traits. In each of the remaining eight scenarios, male MS co-varied positively or negatively with one of the male or female traits described above. The difference in the BG between the null scenario and any covariance scenarios reflects the correlation between MS and RS arising from to confounding (non-causal) effects. This difference represents the extent to which the strength of pre-copulatory sexual selection for mates might be misinterpreted. When simulating covariances between male MS and another trait, we assumed that only the distributions of MS and the trait of interest co-varied. All other (non-covarying) traits were sampled using a Monte-Carlo process from independent, biologically meaningful normal distributions (see Table 1) as follows:

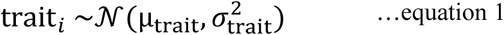

When MS covaried with another trait, these two traits were drawn from a bivariate normal distribution of the form:

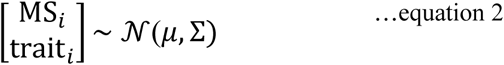

with a mean of 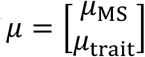 and a variance-covariance matrix given by:

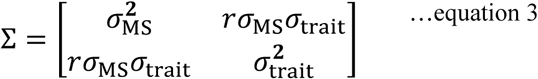

For positive and negative covariances, the correlation coefficient (*r*) was set to +0.707 or - 0.707 respectively (to give an R^2^ of 0.5). In all our simulations, we recorded the order in which females encountered the focal male, and the numbers of offspring the male produced with each female in his mating sequence.

We simulated the following scenarios:

#### Null scenario

In the null scenario (H0), all male and female traits (MS*, Q, V, T, F, P, L*) were sampled independently without any covariances between these traits.

#### Positive covariances due to condition dependence

We simulated two scenarios in which males who are better at acquiring mates also have larger ejaculate reserves (Scenario H1.1: Cov(MS, 𝑄) > 0) or a higher proportion of potent sperm (Scenario H1.2: Cov(MS, 𝑉) > 0). Such positive covariances could occur due to condition dependence, where males in better condition have improved pre- and post- copulatory sexual traits compared to males in poorer condition.

#### Positive covariance due to male-female interactions

We simulated three scenarios in which MS co-varied with female traits. First (Scenario H2.1: Cov(MS, 𝐹) > 0), we created positive covariance between MS and female egg number (*F*). Such a scenario could arise when males who are better at acquiring females also mate with more fecund females. Next (Scenario H2.2: Cov(MS, 𝑃) > 0), we simulated a scenario where MS positively co-varied with female allocation of eggs (*P*), for instance, if males who are better at acquiring females are also better at stimulating female into allocating a higher proportion of eggs for fertilisation. Third (Scenario H2.3: Cov(MS, 𝐿) < 0), we simulated positive covariance between MS and the proportion of the male’s inseminated sperm that the female retains (𝐿). This scenario could arise if females bias sperm retention toward dominant males (who are better at acquiring mates), or if dominant males produce ejaculates that more effectively lead to their sperm being used by the female or entering female sperm storage organs.

Future studies could additionally simulate scenarios where MS negatively co-varies with female polyandry, for example, when males better at acquiring females are also better at suppressing polyandry (or alternatively, males better at acquiring females are more likely to mate with virgin females).

#### Negative covariances due to trade-offs

We simulated negative covariances of male mating success (MS) with male ejaculate size (*Q*), sperm potency (*V*), and the proportion of his stored ejaculate (*T*) that he transfers to a female (Scenarios H3.1: Cov(MS, 𝑄) < 0; H3.2: Cov(MS, 𝑉) < 0; Scenario H4.1: Cov(MS, 𝑇) < 0, respectively). Such scenarios could arise when males, due to resource limitation, face a trade-off between allocating energy towards maintenance/development of pre-copulatory traits (such as ornamentation or weapons) that improve MS, or towards post-copulatory traits such as producing sperm or maintaining sperm quality. These trade-offs often manifest in species with alternative reproductive tactics, in which dominant males that monopolise females have less viable or fewer sperm, or ejaculate smaller proportion of their stored sperm, compared to subordinate males.

### Simulation structure

#### Without sperm competition

The reproductive success (RS_𝑖_) of the *i*^th^ male was the sum of all offspring produced by this male across his mating sequence, and was given by:

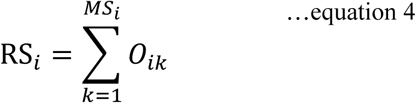

where MS_𝑖_ is the male’s mating success, and *O_ik_* is the number of offspring he produced with the *k*^th^ female he mated with. When there was no sperm competition, the number of offspring produced by the *i*^th^ male with his *k*^th^ mate was calculated as the minimum of: (i) the number of sperm capable of effective fertilisation that the *i*^th^ male transfers to the *k*^th^ female that the female retained (𝑆_𝑖𝑘_), and (ii) the number of eggs capable of being fertilised that the *k*^th^ female allocated to being fertilised by the *i*^th^ male (𝐸_𝑖𝑘_):

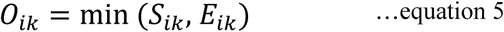

𝑆_𝑖𝑘_ was further partitioned as:

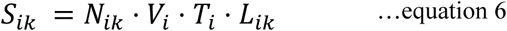

where 𝑁_𝑖𝑘_ was the number of remaining sperm the male had stored in his ejaculate reserves when encountering female *k*^th^, 𝑉_𝑖_ was the proportion of the male’s sperm that were potent/capable of efficient fertilisation, 𝑇_𝑖_ was the proportion of his remaining sperm that the male allocated to the *k*^th^ female, 𝐿_𝑖𝑘_ was the proportion of inseminated sperm that the female retained for fertilisation. The distribution of 𝑆_𝑖𝑘_ values (𝒟_𝑆_) was later used in the models with sperm competition (more below). The number of sperm remaining in the *i*^th^ male’s reserves when encountering the *k*^th^ female (𝑁_𝑖𝑘_) was calculated as his starting ejaculate size before he began mating sequentially with any females (𝑄_𝑖_), minus the number of sperm he had already transferred to the females he copulated with before encountering female *k*.

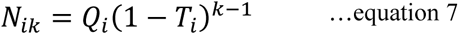

In females, the number of eggs that the *k*^th^ female allocated for being fertilised by the *i*^th^ male, was calculated as the number of eggs this female had when encountering the male(𝐹_𝑖𝑘_) multiplied by the proportion of those eggs which she allocated for being fertilised by the encountered male (𝑃_𝑖𝑘_):

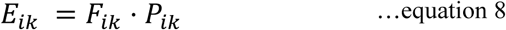

Fisher’s condition says that the total reproductive success of males and females must be equal (Jennions and Fromage, 2017). At equal sex ratios, this means that the mean RS of males and females must be equal. To ensure this, we parametrized the mean of *P* (𝜇_𝑃_) as:

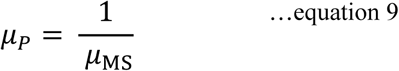

Here, the mean proportion of her eggs that a female allocated to be fertilised by a male (𝜇_𝑃_) equalled the inverse of mean mating success of males (𝜇_MS_) in the population. Our models without sperm competition did not necessarily imply the lack of polyandry, and could also reflect systems where females use sperm from multiple mates sequentially.

#### With sperm competition

Next, we simulated the nine scenarios (H0 to H4.1) in a system where each *k*^th^ female in the *i*^th^ male’s mating sequence re-mated once with another male, right after mating with the *i*^th^ male but before fertilizing any eggs. The number of potent sperm retained by the female from this second, competing male was sampled from the distribution of 𝑆_𝑖𝑘_ values of the focal males from the null scenario without sperm competition (i.e. see equation 6). This distribution of 𝑆_𝑖𝑘_ values (𝒟_𝑆_) represents the number of potent sperm a female retains after mating with a randomly chosen male at a random point in his mating sequence, in the absence of covariances between male mating success (MS) and any other male or female traits (see equation 1). We generated 𝒟_𝑆_ separately for systems with low and high anisogamy (Figure S1). The number of potent sperm from the second male that the female retains (𝐴_𝑖𝑘_) was therefore sampled as:

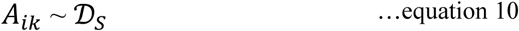

The paternity share of the *i*^th^ male under sperm competition (𝑍_𝑖𝑘_) after his *k*^th^ mate re-mated, was calculated as the proportion of the total potent sperm retained in the female after the two matings, which belonged to the *i*^th^ male.

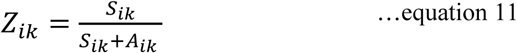

The total number of sperm from both males (*X*) in the female was thus written as:

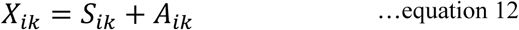

We calculated the number of offspring sired by the *i*^th^ male 𝑂_𝑖𝑘_ with the *k*^th^ female as:

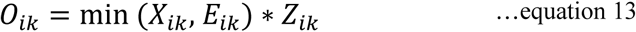

where 𝐸_𝑖𝑘_ is the total number of eggs the female allocates to be fertilised by the two males. To satisfy Fisher’s condition in the sperm competition systems, females allocated twice the number of eggs for fertilisation compared to systems without sperm competition.

### Calculating BG

Bateman gradients (𝛽_𝑠𝑠_) were calculated as the slope (regression coefficient) of the OLS linear regression of male relative reproductive success (𝑟𝑠_𝑖_) on their relative mating success (𝑚𝑠_𝑖_).

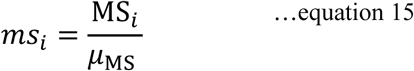

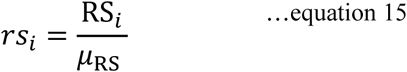

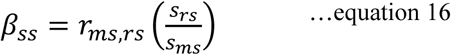

where 𝑟_MS,RS_ is the correlation between *ms* and *rs*, while 𝑠_𝑟𝑠_ and 𝑠_𝑚𝑠_ are their sample standard deviations. In each of our nine simulated scenarios within each of the four biological systems, we simulated 20 replicates with 2000 males per replicate. 𝜇_MS_, 𝜇_RS_, and 𝛽_𝑠𝑠_were calculated separately for each replicate within each scenario and system, and then averaged across the 20 replicates to obtain the final (mean) 𝛽_𝑠𝑠_ for each scenario within each system.

### Simulation assumptions

We made certain assumptions for the sake of simplicity:

- All traits for the focal male and females were sampled from a normal distribution.
- Focal males mated sequentially with females and there was no replenishment of ejaculates.
- Males in the population did not directly interact with each other.
- Males or females did not die during the simulation.
- There was no long term-sperm storage, and the *k^th^* females fertilised her allocated eggs immediately after the *i*^th^ male (in the systems without sperm competition) or his sperm competitor (in systems with sperm competition) had copulated.
- The outcome of sperm competition was determined solely based on the relative proportion of a male’s potent sperm retained in the female. We did not model sperm offence-defence dynamics or precedence.
- Neither males nor females re-mated with the same partner.
- In the system with sperm competition, there was no variance in female degree of polyandry, and all females mated with two males.

### Parametrization (Table 1)

For the low anisogamy system, we parametrized our traits based on data from *Drosophila melanogaster*, because it represents one of the lowest ratios of sperm:egg numbers in animals (Bjork and Pitnick, 2006). In *D. melanogaster*, the observed degree of anisogamy (sperm number reserves in male:egg number reserves in females) is ∼30:1, while it is 6:1 in *D. bifurca* and *D. hydei* (Bjork and Pitnick, 2006; Bjork et al, 2007). We chose 7.5:1 as the degree of low anisogamy for the sake of argument. For the high anisogamy system, we parametrized sperm and egg numbers (thus degree of anisogamy of 100000 sperm:1 egg) based on data from red junglefowl (*Gallus gallus*). The standard deviation (𝜎) for all trait distributions was set as 20% of the mean.

### Data analysis

We simulated nine different scenarios in four biological systems, each across 20 replicates with 2000 males per replicate. We first compared the BG obtained from the null scenario against each covariance scenario within a biological system. The difference in BG between these can be interpreted as the fraction of the BG that does not represent the strength of sexual selection on competition for mates. To do this, we first calculated the *rs* and *ms* for each male, and then 𝛽_𝑠𝑠_, separately for each of the 20 replicates within each scenario and system. Next, we created a linear model with Gaussian error distribution with 𝛽_𝑠𝑠_ as the dependent variable and scenario ID as the fixed effect (20 values per scenario), separately for each of the four systems. The null scenario was set as the reference to allow comparisons of the BG of other covariance scenarios against that of the null scenario. Using ANOVA on the linear model, we additionally tested for overall differences in the means of the different scenarios within each system. We ensured that all our linear models met assumptions of normal distribution of residuals and homoscedasticity. Within each biological system, the distribution of offspring produced with a female through the mating sequence, and of MS, was similar across all scenarios (Figure S2, S3, S4, S5, S6).

Next, we used linear mixed models in the *lme4* package (Bates et al, 2014) to calculate partial Bateman gradients (i.e., BG calculated as partial regression coefficients) that account for linear covariances between MS and other traits (following Henshaw et al, 2018, 2020), and tested whether these partial BGs were similar to BG in the null model. These models were constructed to understand whether confounded relationships between MS and RS can still allow interpretation of the true strength of pre-copulatory sexual selection, if the confounding variable is accounted for. To do this, we only used data from the high anisogamy system with sperm competition because this is arguably the more common of the four systems observed. We fit models for each scenario separately. In a model with Gaussian error distribution, we first fit relative reproductive success (*rs*) as our dependent variable, relative mating success (*ms*) as the only fixed effect, and replicate ID as a random effect (to account for non-independence between different males from the same replicate). We then compared the BG estimated in this model to one where all other traits (*Q, V, T, L, F, P*) as well as *ms* were included as fixed effects. Since the null scenario had no covariances between traits, we predicted that adding these traits would not affect the BG compared to the simpler model with only *ms* as a fixed effect. Next, we created these linear models for each covariance scenario. First, we built a model with only *ms* as a fixed effect; second, a model with *ms* and the co-varying trait as fixed effects; and third, a model with *ms* and all traits (*Q, V, T, L, F, P*) as fixed effects. We predicted that the model with only MS as a fixed effect would produce an inflated or deflated BG compared to the BG of the null model, due to unaccounted covariances. In the second model, accounting for the co-varying trait would give a partial BG similar to that measured in the null scenario. In the third model, adding non-co- varying traits would not further alter the BG compared to the second model.

## Results

### Low anisogamy, no sperm competition

Positive covariances between male mating success (MS) and *Q* (starting ejaculate size)*, V* (proportion of potent sperm)*, F* (number of eggs in female)*, P* (proportion of allocated eggs)*, or L* (proportion of sperm retained) led to significantly steeper BG, while negative covariances between MS with *Q* or *V* resulted in significantly shallower BG (Figure 2, Table S1, Figure S7, S8), compared to the null scenario. Negative covariance between MS and *T* (proportion of stored sperm transferred when copulating) produced a significantly steeper BG than the null scenario.

**Figure 2:**
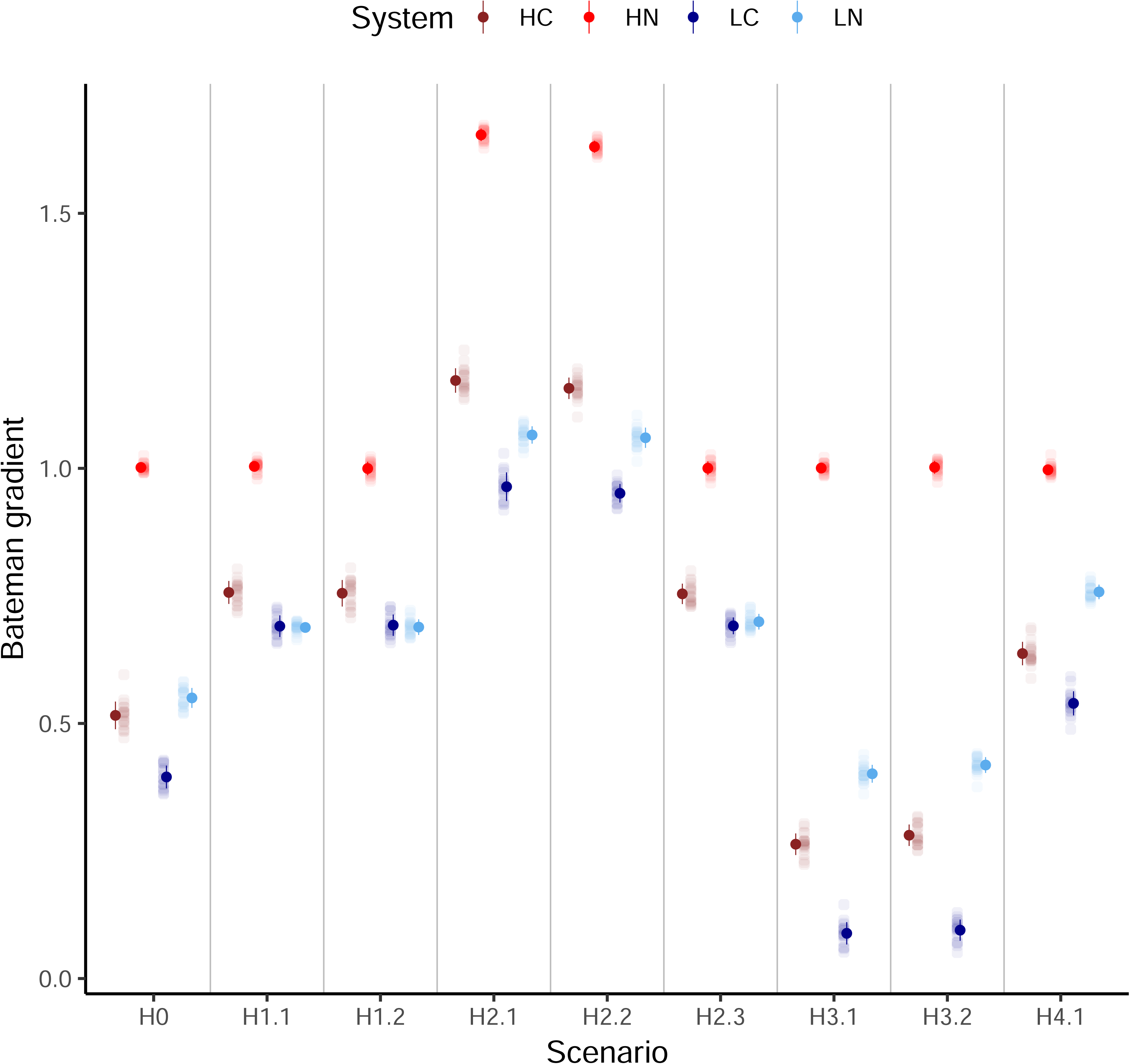
Bateman gradients for each of the nine scenarios across the four biological systems. Y axis represents the average increase in *rs* when *ms* increases by one unit. Dots show mean BG per scenario per system, error bars show standard deviations of the BG across 20 replicates (jittered). **HC:** High anisogamy with sperm competition; **HN:** High anisogamy without sperm competition; **LC:** Low anisogamy with sperm competition; **LN:** Low anisogamy without sperm competition. H0: no covariances; H1.1: positive covariance between MS and ejaculate size (*Q*); H1.2: positive covariance between MS and sperm potency (*V*); H2.1: positive covariance between MS and female egg number (*F*); H2.2: positive covariance between MS and female egg allocation (*P*); H2.3: positive covariance between MS and female sperm retention (*L*); H3.1: negative covariance between MS and ejaculate size (*Q*); H3.2: negative covariance between MS and sperm potency (*V*); H4.1: negative covariance between MS and proportion of stored ejaculate transferred by male to female (*T*). See Figures S7 and S8 for BGs represented using regressions.

Visual inspection (Figure 3) of offspring production through the mating sequence revealed that in the null scenario (H0), patterns of offspring production were the same for males with different MS. In scenarios where MS co-varied positively with *Q*, *V*, or *L* (H1.1, H1.2, H2.3), males with higher MS had shallower rates of decline in offspring production compared to males with lower MS. Conversely, in scenarios where MS co-varied negatively with *Q* or *V* (H3.1, H3.2), males with higher MS experienced steeper declines in offspring production through the mating sequence. When MS and *T* negatively co-varied (H4.1), males with lower MS experienced steeper declines in offspring production through the mating sequence compared to males with higher MS. When MS co-varied positively with *F* or *P*, males with higher MS consistently produced more offspring than males with lower MS, irrespective of female rank (H2.1, H2.2).

**Figure 3:**
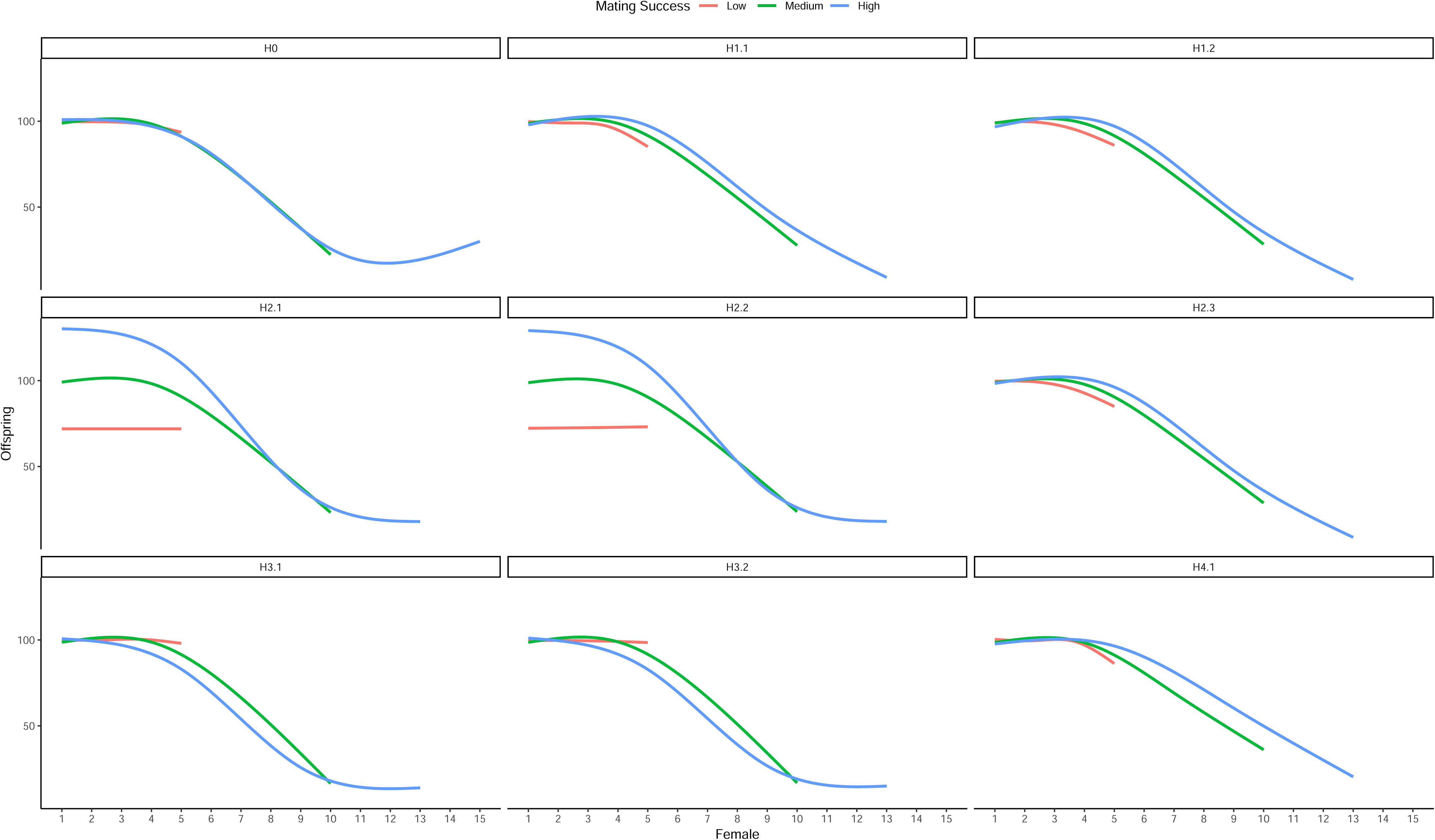
Effect of MS on the number of offspring a focal male produces with each female (rank) in his mating sequence. Panel labels correspond to each scenario in the system with low anisogamy and no sperm competition. Data combined from all 20 replicates. MS binned into three categories for ease of visualisation. Lines show means of 20 replicates, constructed as gam smooths with four knots in ggplot. H0: no covariances; H1.1: positive covariance between MS and ejaculate size (*Q*); H1.2: positive covariance between MS and sperm potency (*V*); H2.1: positive covariance between MS and female egg number (*F*); H2.2: positive covariance between MS and female egg allocation (*P*); H2.3: positive covariance between MS and female sperm retention (*L*); H3.1: negative covariance between MS and ejaculate size (*Q*); H3.2: negative covariance between MS and sperm potency (*V*); H4.1: negative covariance between MS and proportion of stored ejaculate transferred by male to female (*T*).

### Low anisogamy, with sperm competition

Positive covariances between MS and *Q, V, L, F,* or *P* led to significantly steeper BG than the null scenario (Figure 2, Table S2, Figure S7, S8), while negative covariances of *MS* with *Q* or *V* produced a shallow BG. Negative covariance between MS and *T* resulted in a significantly steeper BG than the null scenario (Table S2). Patterns of offspring production across the mating sequence were the same for focal males with different MS in the null scenario (Figure 4). However, when male MS co-varied positively with *Q*, *V*, *L*, *F* or *P*, males with higher MS consistently produced more offspring with each female compared to males with lower MS, irrespective of female rank in the mating sequence. On the other hand, when MS co-varied negatively with *Q* or *V*, males with higher MS consistently produced fewer offspring with each female irrespective of female rank, compared to males with lower MS. Additionally, when MS co-varied positively with *T*, males with lower MS produced more offspring with females early in the sequence, but fewer offspring with females later in the sequence, compared to males with higher MS.

**Figure 4:**
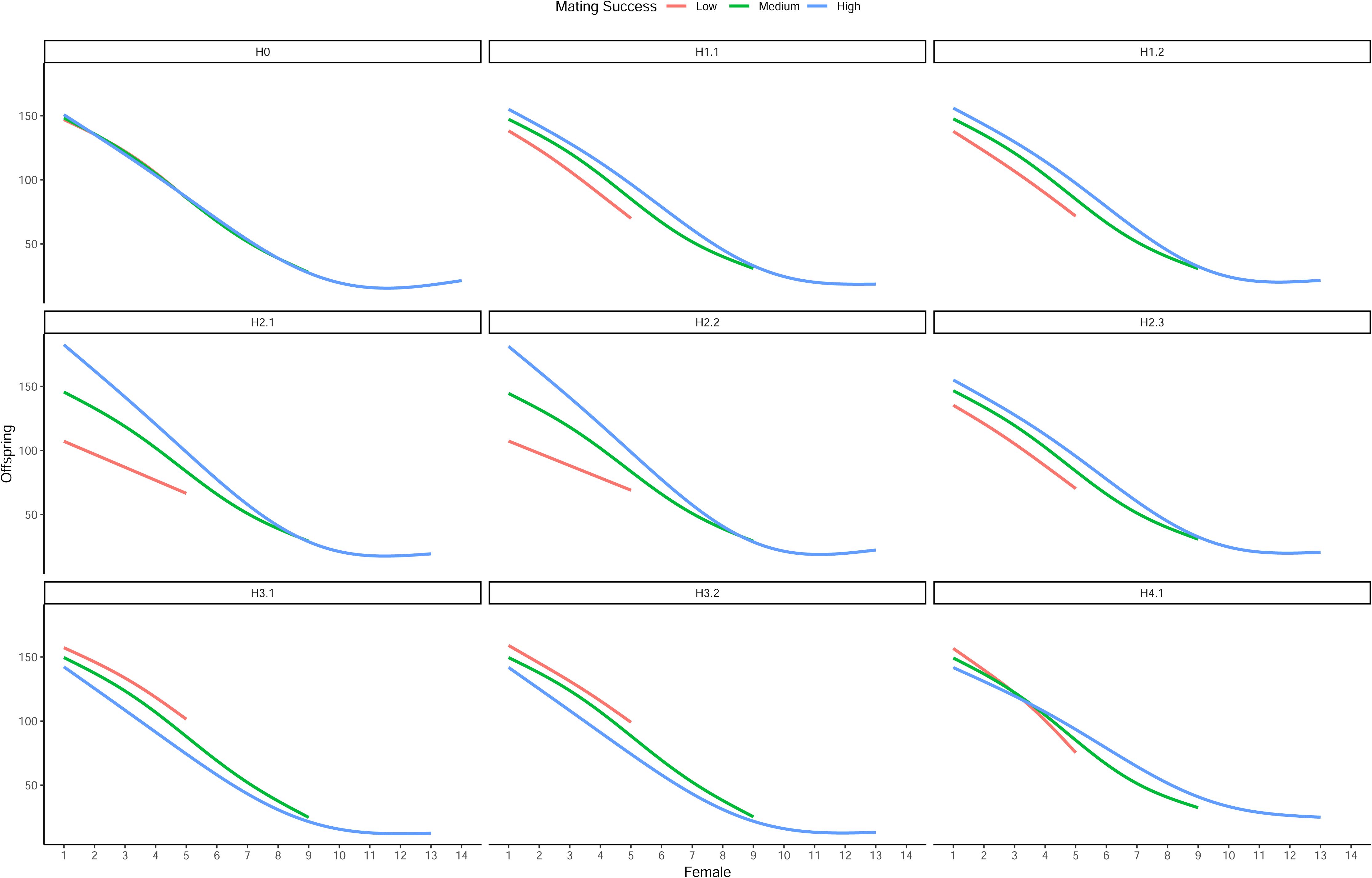
Effect of MS on the number of offspring a focal male produces with each female (rank) in his mating sequence. Panel labels correspond to each scenario in the system with low anisogamy and sperm competition. MS binned into three categories for ease of visualisation. Lines show means of 20 replicates, constructed as gam smooths with four knots in ggplot. H0: no covariances; H1.1: positive covariance between MS and ejaculate size (*Q*); H1.2: positive covariance between MS and sperm potency (*V*); H2.1: positive covariance between MS and female egg number (*F*); H2.2: positive covariance between MS and female egg allocation (*P*); H2.3: positive covariance between MS and female sperm retention (*L*); H3.1: negative covariance between MS and ejaculate size (*Q*); H3.2: negative covariance between MS and sperm potency (*V*); H4.1: negative covariance between MS and proportion of stored ejaculate transferred by male to female (*T*).

### High anisogamy, no sperm competition

Positive covariances only between MS and *F* or *P*, led to significantly steeper BG than the null scenario (Figure 2, Table S3, Figure S7, S8). However, BG from the null scenario did not significantly differ from BGs of scenarios in which MS co-varied with *Q*, *V*, *T*, or *L* (Table S3). Patterns of offspring production through the mating sequence were modulated by MS only when MS co-varied positively with *F* or *P* (Figure 5). Here, males with higher mating success consistently produced more offspring than males with lower MS, irrespective of female rank. In contrast to the systems with low anisogamy, offspring production of focal males did not decline through the mating sequence in the high anisogamy system without sperm competition.

**Figure 5:**
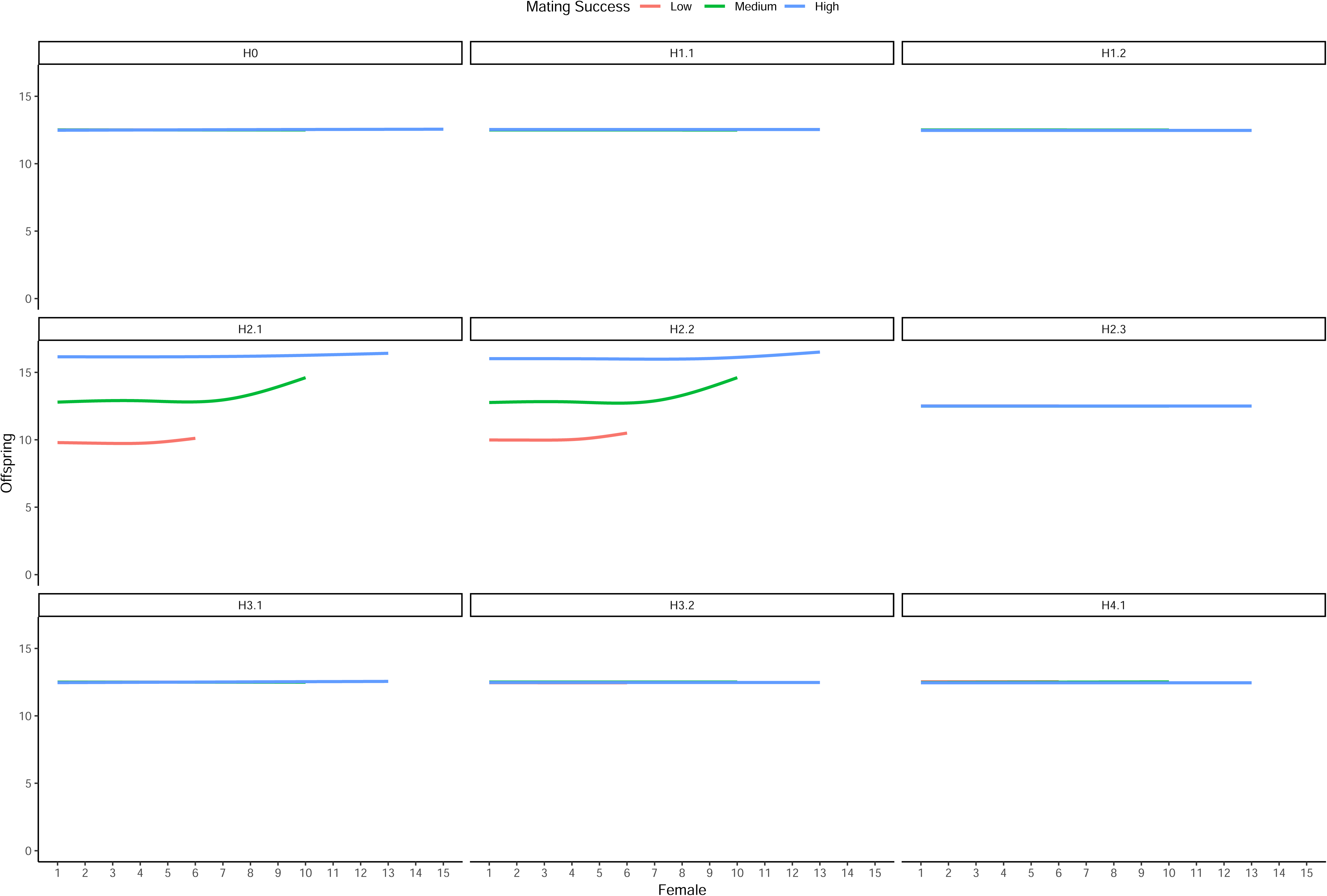
Effect of MS on the number of offspring a focal male produces with each female (rank) in his mating sequence. Panel labels correspond to each scenario in the system with high anisogamy and no sperm competition. MS binned into three categories for ease of visualisation. Lines show means of 20 replicates, constructed as gam smooths with four knots in ggplot. H0: no covariances; H1.1: positive covariance between MS and ejaculate size (*Q*); H1.2: positive covariance between MS and sperm potency (*V*); H2.1: positive covariance between MS and female egg number (*F*); H2.2: positive covariance between MS and female egg allocation (*P*); H2.3: positive covariance between MS and female sperm retention (*L*); H3.1: negative covariance between MS and ejaculate size (*Q*); H3.2: negative covariance between MS and sperm potency (*V*); H4.1: negative covariance between MS and proportion of stored ejaculate transferred by male to female (*T*). The increase in offspring production with later females is not a true increase, but is a consequence of each binned category being an average of multiple mating success values.

### High anisogamy with sperm competition

Positive covariances between MS and ejaculate size (*Q*, H1.1), sperm potency (*V*, H1.2), sperm retained by females (*L,* H2.3), female egg numbers (*F*, H2.1) or egg allocation (*P,* H2.2), led to significantly steeper BG, while negative covariance between male MS and *Q* (H3.1) or *V* (H3.2) led to a significantly shallower BG, than the null scenario (Figure 2, Table S4, Figure S7, S8). Negative covariance between MS and the proportion of a male’s stored sperm transferred to a female (*T*, H4.1), led to significantly steeper BG than the null scenario. Offspring production through the mating sequence was not modulated by MS in the null scenario. However, in all covariance scenarios, MS modulated offspring production (Figure S9), and did so in a similar way as in the low anisogamy system with sperm competition.

We also evaluated whether adding the co-varying trait in each covariance scenario as a fixed effect, led to the estimated BG being closer to the null scenario, thus better representing the causal influence of MS on RS (Figure 6). In the null scenario where no covariances existed, including additional traits did not alter the influence of MS on RS. However, in scenarios where MS co-varied with another trait, the partial BG obtained by including the co-varying trait as a linear fixed effect, led to substantially similar estimates to the BG from the null scenario, compared to models that did not include the co-varying trait (Figure 6).

**Figure 6:**
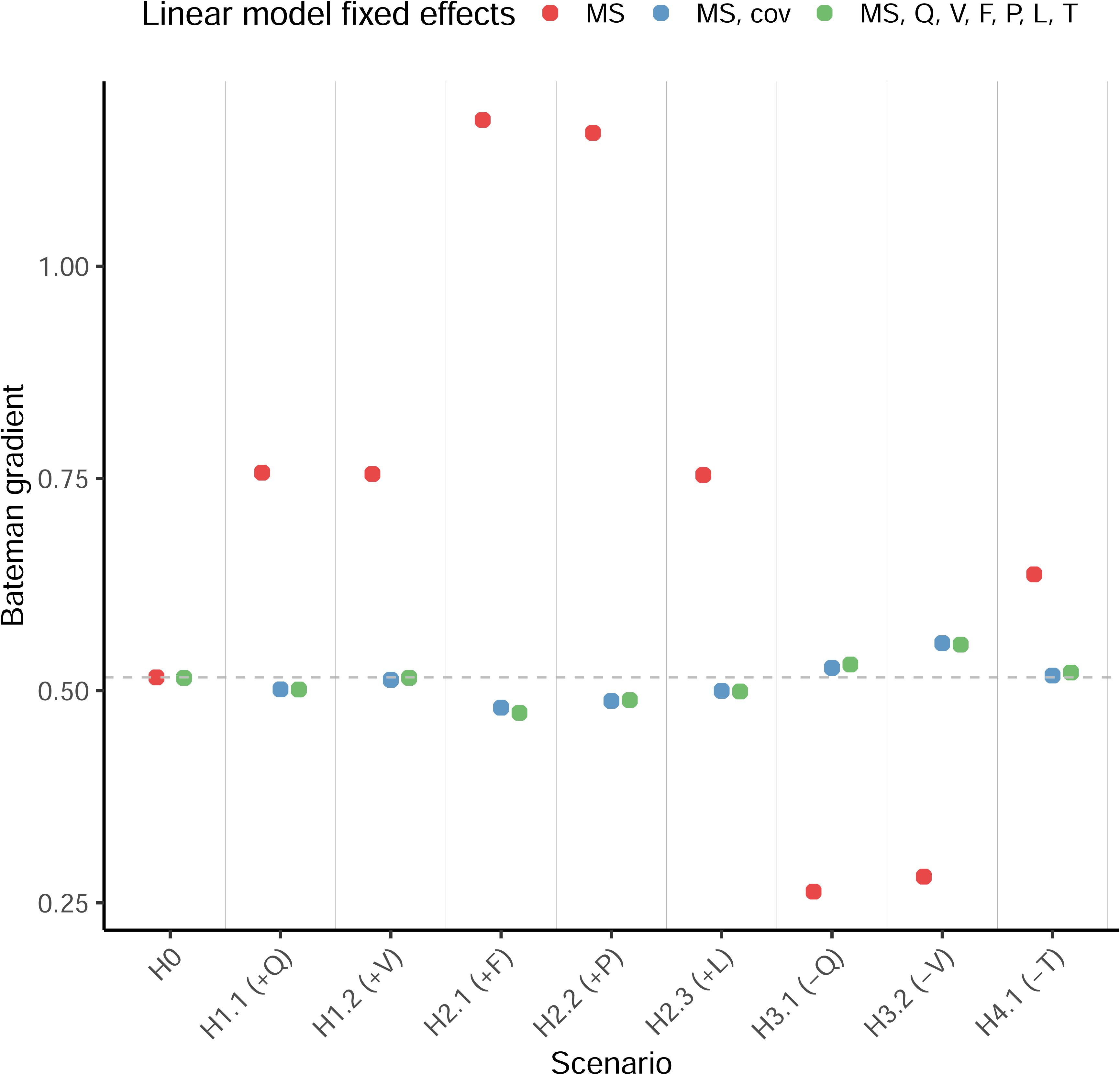
Comparison of Bateman gradients estimated through linear models across nine different scenarios, in the system with high anisogamy and sperm competition, with or without accounting for covariances. A possible reason why the partial BG in each covariance scenarios is not exactly the same as the BG in the null scenario, is because the covariates are modelled as linear terms, however, their influence on RS may be non-linear. Additionally, random sampling differences of traits between the scenarios might lead to “true” variation in the BG between scenarios. In the null scenario (H0), there are no covariances between MS and any other traits (*Q, V, F, P, L, T*). In other scenarios (H1.1 to H4.1) the specific trait that co-varies (cov) with MS is specified in parentheses, along with the direction of covariance (positive “+” or negative “–“). In red, BG from linear models where only *ms* is included as a fixed effect, with *rs* as the dependent variable. In blue, partial BG from models where only the co-varying variable (cov) for that scenario along with *ms* are included as fixed effects. In green, partial BG from linear models where all traits (including the co-varying variable, if any) and *ms* are included as fixed effects. Mean estimates of the BG from the linear models shown as dots (N = 2000 males per scenario across 20 replicates). Dotted horizontal line shows the BG for the null scenario.

## Discussion

We conducted simulations to explore how male mating success (MS) correlates with reproductive success (RS) due to both, the causal effect of MS on RS, and covariances between MS and other male and female reproductive traits. The Bateman gradient (BG) is typically used to interpret the strength of pre-copulatory sexual selection, i.e. the potential improvement in fitness obtained by acquiring more mates (Wade & Shuster, 2005). Males, due to having steeper BGs, are typically hypothesized to experience more intense pre-copulatory sexual selection than females, leading to predictions of males competing for mates and being promiscuous, more than females. However, consistent with previous critiques (Anthes et al, 2010, 2017; Henshaw et al, 2018; Tang-Martinez, 2016), we show that without accounting for covariances, the measured BG can be misleading. This is because under covariances, the BG reflects not just the strength of sexual selection on intra-sexual competition for mates, but also the strength of other forms of selection, such as post-copulatory sexual selection on traits like ejaculate size and fertilization efficiency of sperm (Anthes et al, 2010, 2017; Pischedda and Rice, 2012). Modulation of the BG due to such covariances has important implications for quantifying sex-differences in sexual selection. For instance, if a steeper BG in males than females (as typically predicted) in a species are caused by covariances between male MS and ejaculate traits rather than a causal effect on RS, then the difference in BGs between sexes cannot be interpreted as stronger pre-copulatory sexual selection on males than females.

A fundamental puzzle in sexual selection is why male BGs vary within (e.g. *Drosophila melanogaster*, reviewed in: Davies et al, 2023) as well as between species (Anthes et al, 2017). Our simulations provide some explanations for inter- and intra-specific variation in BG, solely based on the presence or absence of some covariances in certain populations. For instance, in populations where males vary in their condition, high condition males might have more competitive ejaculates as well as higher MS, compared to low condition males (Puniamoorthy et al, 2012; Reznick et al, 2000). Such covariances due to condition dependence (e.g. Hare and Simmons, 2022; Janicke et al, 2015; Mehlis et al, 2015; Morimoto et al, 2016) would lead to the measured BG being steeper in these populations compared to populations of the same species where male condition is more homogeneous. On the other hand, in species with alternative mating tactics, negative covariances between male MS and ejaculate size, quality, or allocation (e.g. Alonzo and Warner, 2000; Durant et al, 2016; Froman et al, 2002; Gage et al, 1995; Klaus et al, 2011; Lupold et al, 2014; Preston et al, 2001; Saunders and Shuster, 2020; Warner et al, 1995) could lead to a shallower male BG compared to species without such tactics. For instance, males in dominant roles in species with alternative mating tactics often achieve similar overall reproductive success as males in subordinate roles (Da Silva et al, 2024). In such scenarios, the relationship between MS and RS could be curvilinear (see Figure S8) with male fitness being maximised at average MS. The BG might also be inflated in species with condition-based assortative mating, where males in high condition (who have high MS) mate with females in high condition (who have high fecundity) (e.g. Collet et al, 2014; Gerlach et al, 2012; Greenway et al, 2021; Kraak and Bakker, 1998). Importantly, when populations or species differ in their BGs due to the presence of certain covariances, the BGs cannot be interpreted as different strengths of pre-copulatory sexual selection on acquiring mates between these populations/species.

Variation in BGs between different species could also be due to differences in anisogamy or sperm competition between taxa. Anisogamy underpins the evolution of divergent reproductive strategies, and an increase in the degree of anisogamy is predicted to lead to a steeper BG for the sex with more numerous gametes (Henshaw et al, 2022; Janicke, 2024; Jones et al., 2002; Lehtonen, 2022; Lehtonen & Parker, 2024). Our findings suggest that in the absence of sperm competition, males in high anisogamy systems indeed gain more fitness benefits by improving their MS, compared to males in low anisogamy systems (Figure 2). However, if sperm competition is present within high anisogamy systems, the differences in BG between high and low anisogamy systems becomes less apparent (Figure 2). Our findings generally agree with Arnold (1994), who suggested that a positive, linear relationship between MS and RS for males can only be expected when their RS is not limited by gamete production rate (e.g. in high anisogamy systems without sperm competition).

The importance of partitioning the measured BG into episodes of selection on pre- versus post-copulatory traits has been previously emphasized (Anthes et al, 2017; Evans and Garcia-Gonzalez, 2016; Pelissie et al, 2014). By including post-copulatory processes such as cryptic female choice via differential sperm retention or female fecundity stimulation (see Eberhard, 2015; Firman et al, 2017), and their covariances with MS, our simulations highlight the under-appreciated role of female agency in impacting male BG. Our results further show that it is difficult to disentangle pre- from post- copulatory episodes of sexual selection when using the BG, a phenomenological (rather than process-based) metric.

While our simulations demonstrate that covariances between MS and other ejaculate or female traits can severely distort interpretation of the strength of sexual selection, this does not diminish utility of the BG. We suggest methods for researchers to assess and account for covariances when calculating the BG. First, visually examining offspring production of males with each female in the mating sequence can allow identification of the presence, and importantly, the type of underlying covariances (see Figures 3, 4, 5, 6). In the absence of covariances, no systematic differences should exist in the rate of change or the intercept for offspring production across the mating sequence, between males with varying mating success (see Sanghvi et al, 2024a who show this). When change in offspring production through the mating sequence is linear, the presence of covariances can further be statistically tested for by fitting:

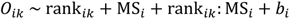

where 𝑂_𝑖𝑘_ is the number of offspring produced by each focal male mated with each female, rank is the rank/order of the female encountering the male in his mating sequence, and 𝑏_𝑖_ is the random effect of male ID to control for male-level non-independence in offspring production. Here, a significant effect of 𝑟𝑎𝑛𝑘_𝑖𝑘_: MS_𝑖_ or of MS_𝑖_ might suggest presence of covariances between MS and other variables that indirectly influence male RS. Second, we suggest that studies include the co-varying trait in the linear model when calculating *β_ss_*, and interpret the strength of pre-copulatory sexual selection only using the partial BG (following Henshaw et al, 2018). While Anthes et al (2017) caution against using this method, we show that in the absence of better alternatives, this approach provides a reasonable approximation of the causal effect of manipulating MS on RS. When the influence of the co-varying trait on RS is linear, the partial BG would represent the causal influence of MS on RS.

Fitness is not only determined by the number of offspring produced, but also by the reproductive value of each offspring (Gillespie et al, 2008). Therefore, the strength of pre-copulatory sexual selection on acquiring mates will be determined by the influence of MS on RS, as well as of MS on offspring quality (Kokko and Jennions, 2008). This distinction is crucial when MS influences offspring quality and RS in contrasting ways (Arnold, 1994; Queller, 1997). For example, when males who mate with more females produce more offspring but are less able to provision each offspring thereby reducing offspring quality (e.g. Lynch, 2016), males with average mating success would likely produce the most numbers of grand offspring despite producing fewer offspring than males with the highest MS (reviewed in Stiver and Alonzo, 2009). Here, despite the BG being positive and the relationship between MS and RS being linear, interpreting the BG as evidence of strong sexual selection on mate acquisition would be misleading. This is because selection is likely to favour a strategy that maximizes the production of grand offspring produced rather than merely of offspring (i.e. RS). Similarly in females, polyandry (i.e. increasing female MS) could confuse the paternity of males, leading to various males provisioning a female’s offspring, which consequently improves offspring quality (Hosken and Stockley, 2003). Here, despite polyandry not directly improving female RS and the BG being shallow, polyandry might still lead to increased numbers of grand offspring via improved offspring quality (reviewed in Jennions and Petrie, 2000; Parker and Tang-Martinez, 2005; Simmons, 2005). In such a case, interpreting a shallow female BG as weak pre-copulatory sexual selection would be misleading. We therefore recommend that covariances between MS and offspring quality, in addition to those between MS and RS, be considered within the BG framework.

Our simulations were limited in their complexity because we did not manipulate degree of polyandry, sperm precedence, male and female lifespans, sperm storage duration, the rate of ejaculate and egg replenishment, offspring survival, or offspring reproductive success. Future studies could test how covariances between MS and these traits might influence the BG and the strength of pre-copulatory sexual selection. For instance, if males who are more attractive (and thus have higher MS) also possess genes that improve offspring survival until detection (e.g. Head et al, 2005), the BG would be modulated. Our simulations focussed only on male BG. However, female BG can also be influenced by covariances between RS and other male or female traits (e.g. Collett et al, 2014; Fromonteil et al, 2023; Gerlach et al, 2012). Future studies could simulate various male- and female-specific covariances in polygynandrous systems to test whether sex-differences in the strength of sexual selection become less apparent after accounting for such covariances (by using partial BGs). Finally, we emphasize that studies which experimentally manipulate MS are alarmingly lacking and encourage researchers to compare BGs obtained from experimental manipulation of MS (e.g. Andrade and Kasumovic, 2005) versus from spontaneously available variation in MS, to understand how covariances might modulate the BG in nature.

## Conclusions

Our study adds to the growing body of research advocating for a more nuanced understanding and application of the Bateman Gradient (Anthes et al, 2017; Parker & Tang-Martinez, 2005; Henshaw et al, 2016). We illustrate that BG measurements which fail to account for covariances between MS and various other traits, can oversimplify its interpretation and misattribute the selection agent to solely being pre-copulatory competition for mates. However, under covariances, the BG partly represents selection on various ejaculate traits and post-copulatory processes. The impact of these co-varying traits depends on the biology of the system, such as the degree of anisogamy and its interaction with level of sperm competition, which could partly explain taxonomic variation in the BG. Overall, we use specific biological examples to demonstrate the range of scenarios where BGs are likely to be misleading; quantify the magnitude and direction of how covariances might modulate the BG (thus the degree by which sexual selection might be misinterpreted when using the BG); reveal how fundamental biological differences between species modulate the BG; and suggest methods by which studies can assess the presence, magnitude, and type of covariances as well as account for them to better interpret the BG. Our simulations offer generalisable insights into the mechanisms that cause variation in the BG, improve our understanding of what the BG represents and what it does not, as well as provide novel insights into sexual selection.

## Supporting information

Supplementary figures (S1 -S10) and supplementary tables (S1-S4) are provided along with this manuscript.

## Acknowledgements

We thank members of the fly lab for their helpful comments. KS is very grateful to Hanna Kokko, Tommaso Pizzari, and Geoff Parker for their suggestions at early stages.

## Author contributions

**KS**: conceptualisation, formal analysis, investigation, writing (original draft); **AK, JMH, TJ**: conceptualisation, writing (review and editing); **IS**: conceptualisation, supervision, writing (review and editing), funding acquisition.

## Data availability

Simulation code and simulated data can be found at Open science framework DOI:10.17605/OSF.IO/WPE8M.

## Abbreviations

MS: mating success
RS: reproductive success
BG: Bateman gradient

## Supporting information

### Supplementary Figures

**Figure S1:**
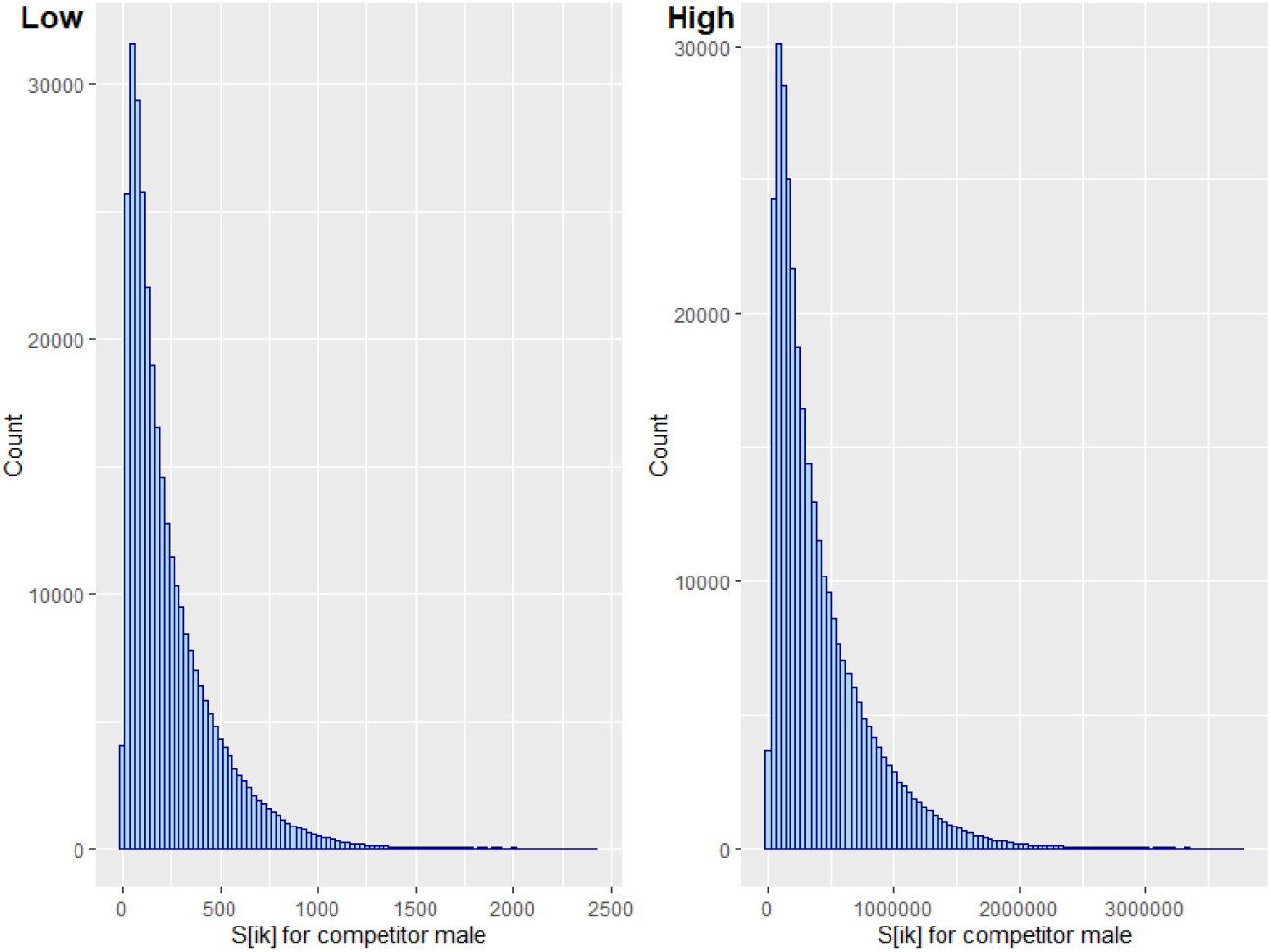
Distribution of *S_ik_* values for the competitor male in low and high anisogamy systems, from which *A_ik_* values for the competitor male’s sperm are sampled. *S_ik_* represents the numbers of potent sperm transferred by a focal male and retained by a mated female, if she mated with a randomly chosen focal male at a random point in his mating sequence.

**Figure S2:**
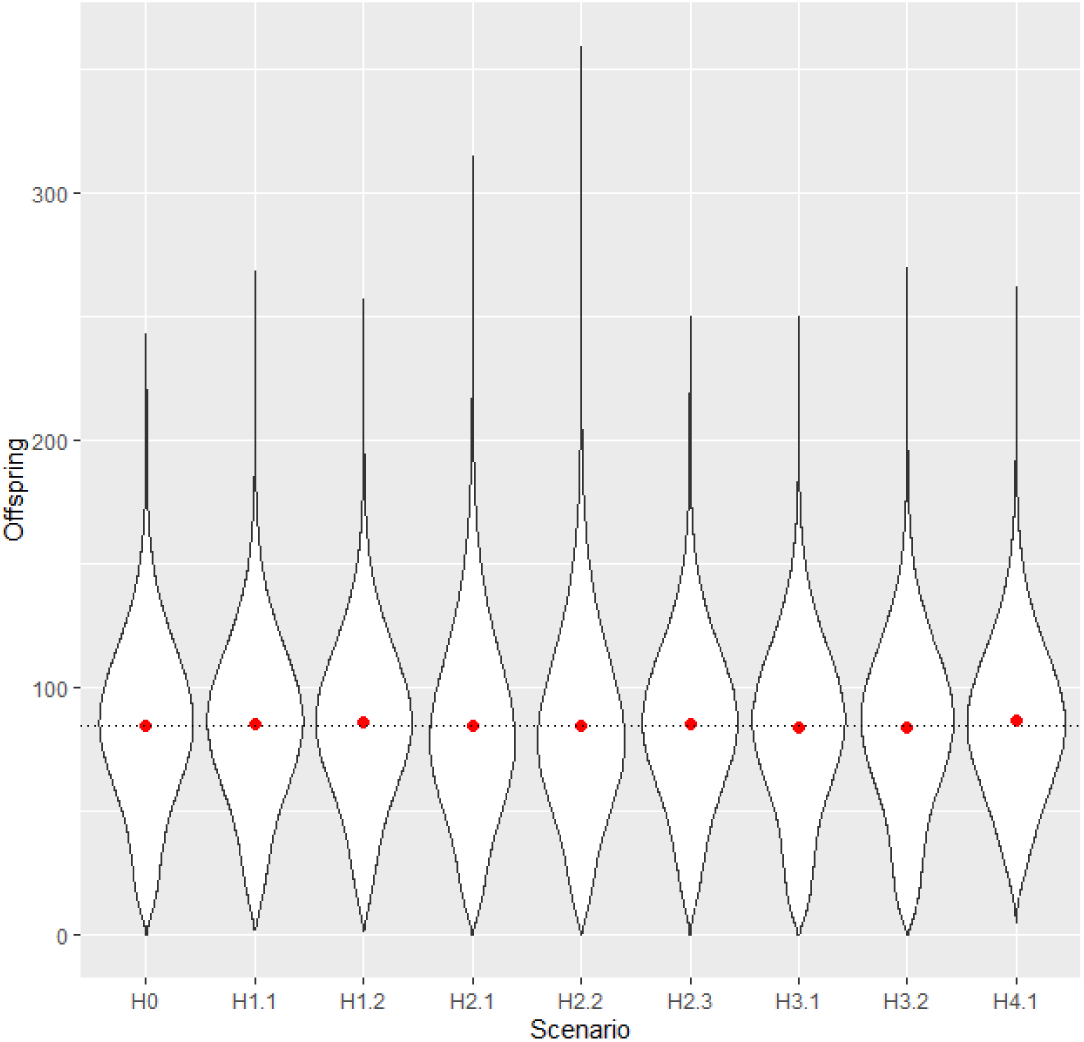
Distribution of offspring production of the focal male with each mated female, across the nine simulated scenarios, in the system with low anisogamy and no sperm competition. Offspring production was similar across scenarios, therefore differences in BG across scenarios cannot be attributed to difference in mean offspring production. Violin plots represent distribution of offspring produced by each focal male with each female, red dot represents mean offspring number per scenario. Dotted line shows overall mean across all scenarios. Data pooled from 20 replicates.

**Figure S3:**
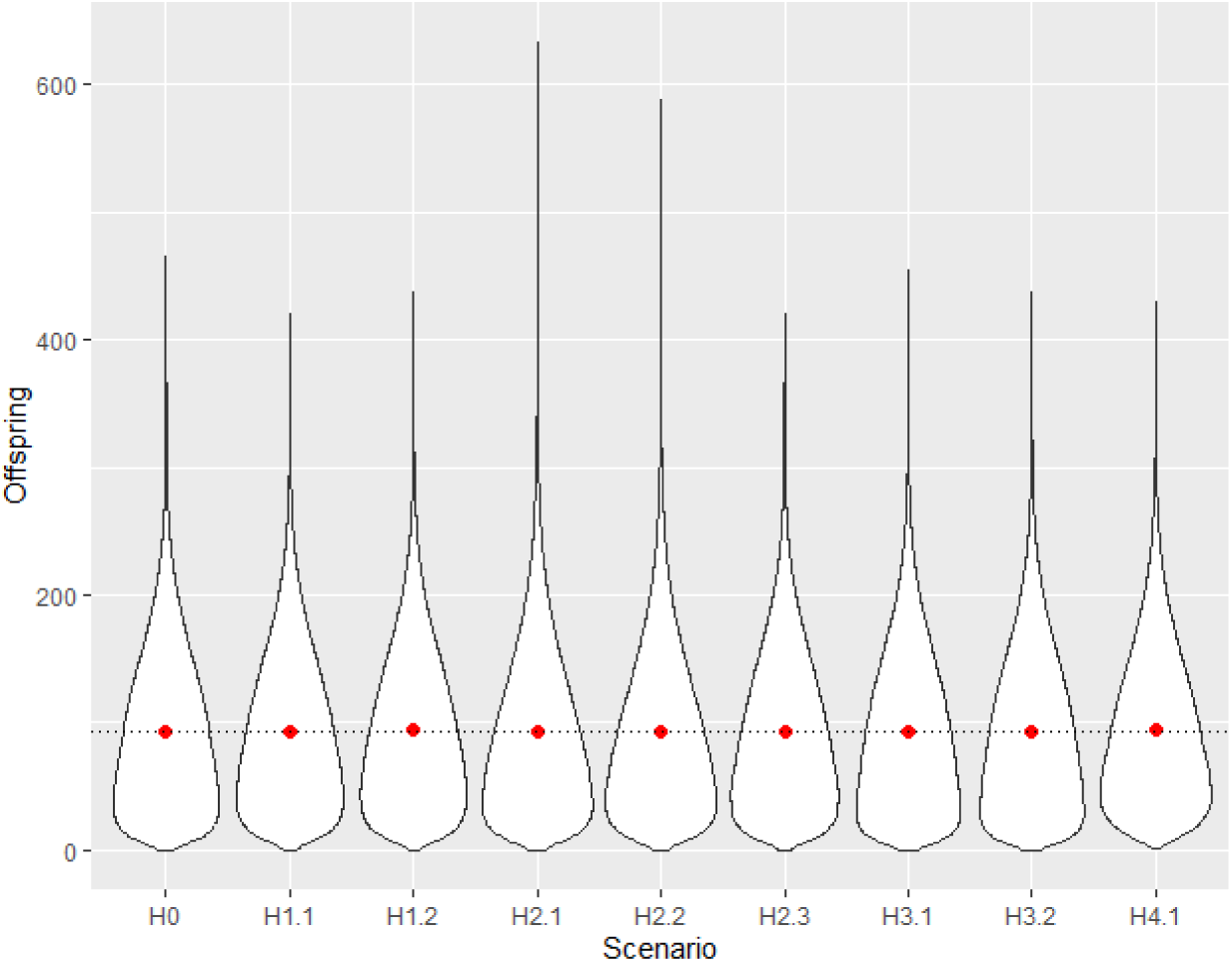
Distribution of offspring production of the focal male with each mated female, across the nine simulated scenarios, in the system with low anisogamy with sperm competition. Offspring production was similar across scenarios, therefore differences in BG across scenarios cannot be attributed to difference in mean offspring production. Violin plots represent distribution of offspring produced by each focal male with each female, red dot represents mean offspring number per scenario. Dotted line shows overall mean across all scenarios. Data pooled from 20 replicates.

**Figure S4:**
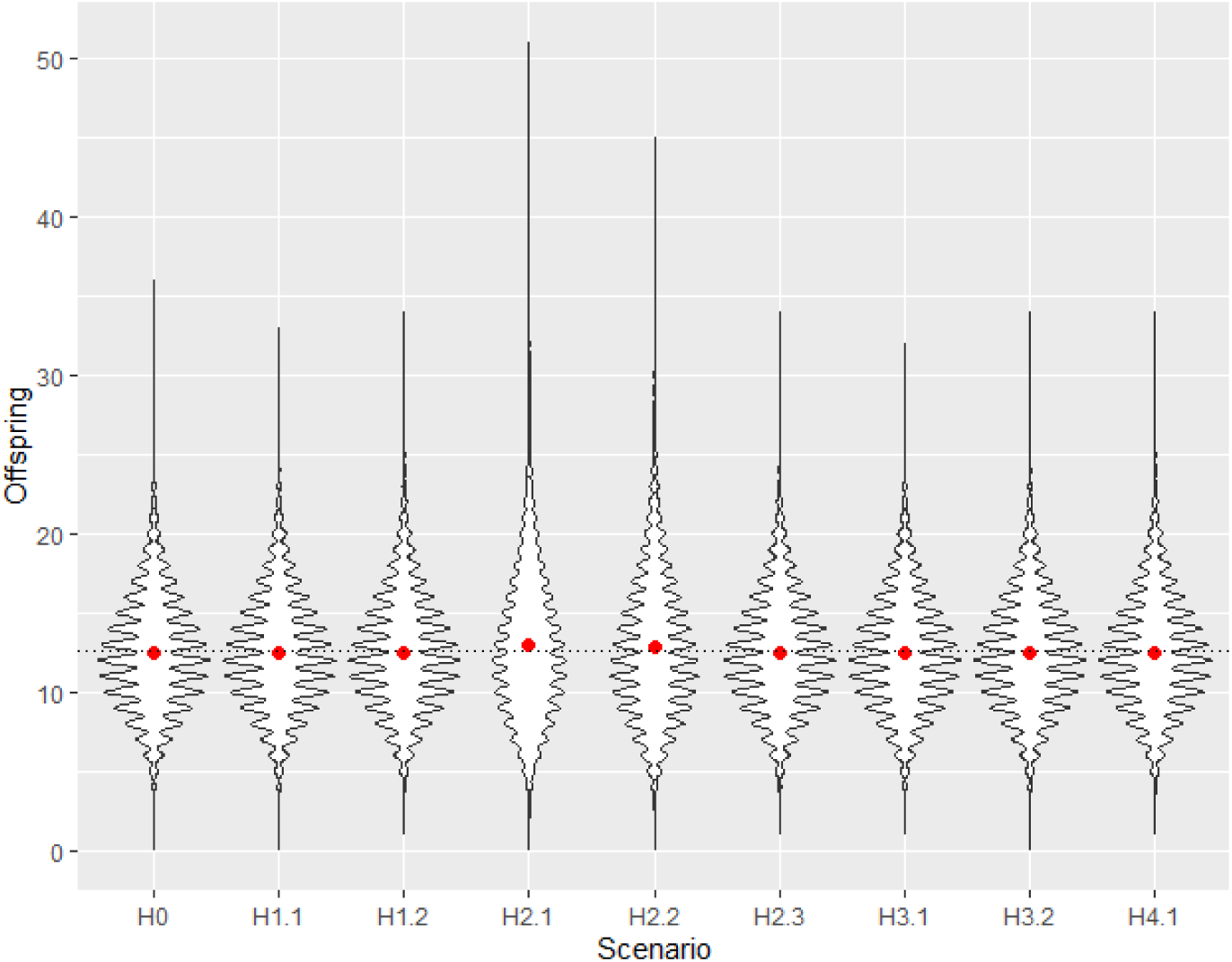
Distribution of offspring production of the focal male with each mated female, across the nine simulated scenarios, in the high anisogamy system without sperm competition. Offspring production was similar across scenarios, therefore differences in BG across scenarios cannot be attributed to difference in mean offspring production. Violin plots represent distribution of offspring produced by each focal male with each female, yellow triangle represents mean offspring number per scenario. Dotted line shows overall mean across all scenarios. Data pooled from 20 replicates.

**Figure S5:**
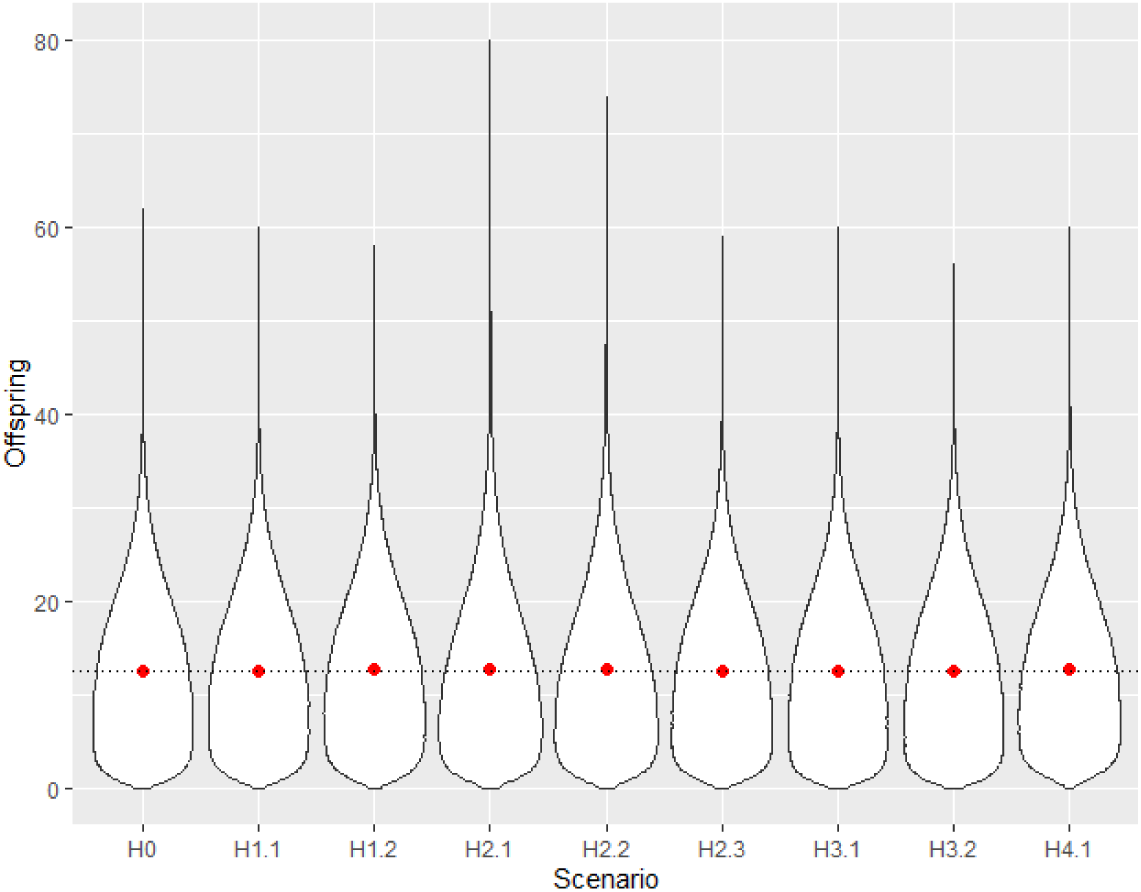
Distribution of offspring production of the focal male with each mated female, across the nine simulated scenarios, in the high anisogamy system with sperm competition. Offspring production was similar across scenarios, therefore differences in BG across scenarios cannot be attributed to difference in mean offspring production. Violin plots represent distribution of offspring produced by each focal male with each female, red dot represents mean offspring number per scenario. Dotted line shows overall mean across all scenarios. Data pooled from 20 replicates.

**Figure S6:**
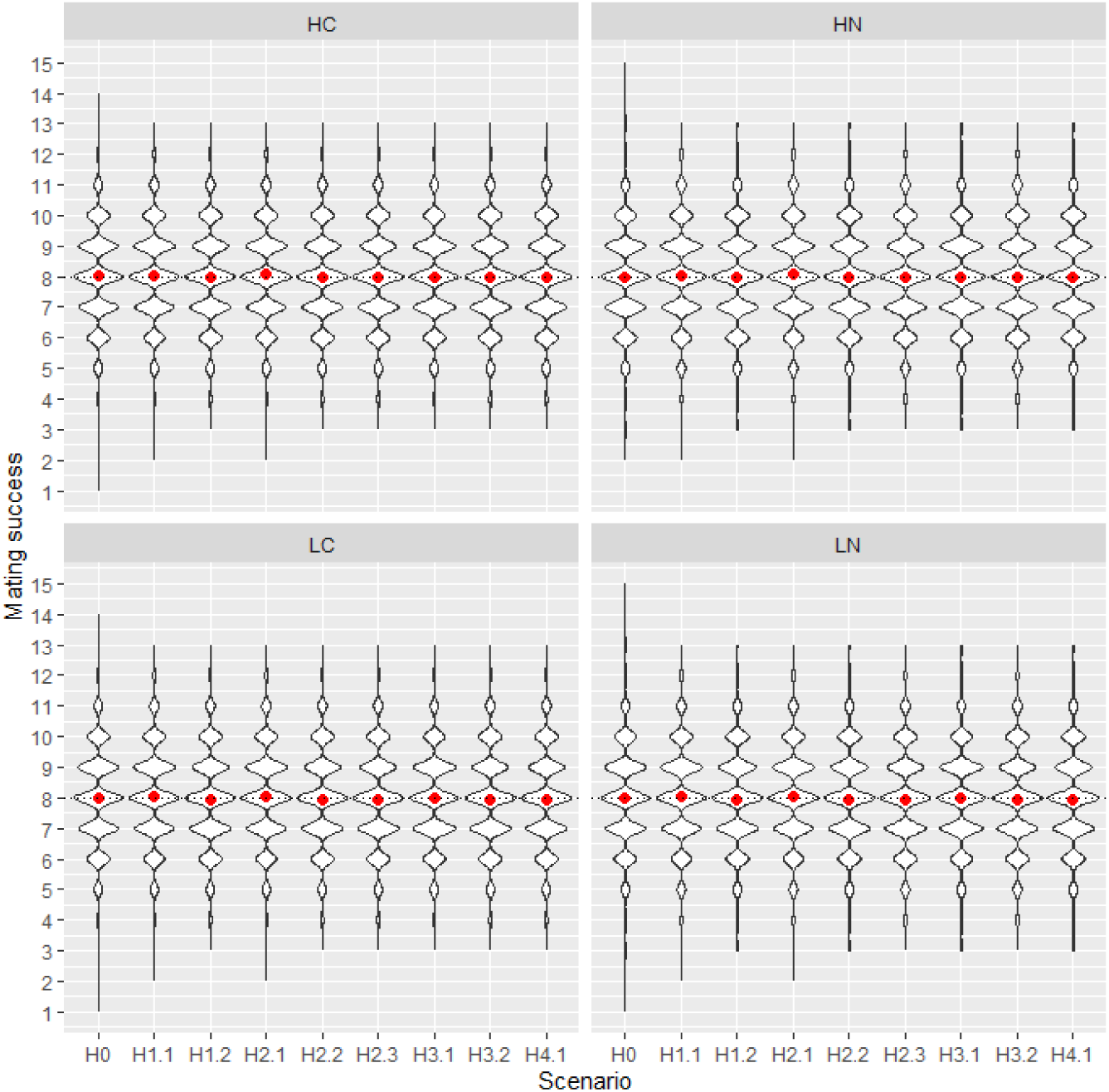
Distribution of male mating success across the nine scenarios and 4 systems. Note that no males in our simulations had a mating success of <1 or >15. High (H) or low anisogamy (L), with (C) or without (N) sperm competition.

**Figure S7:**
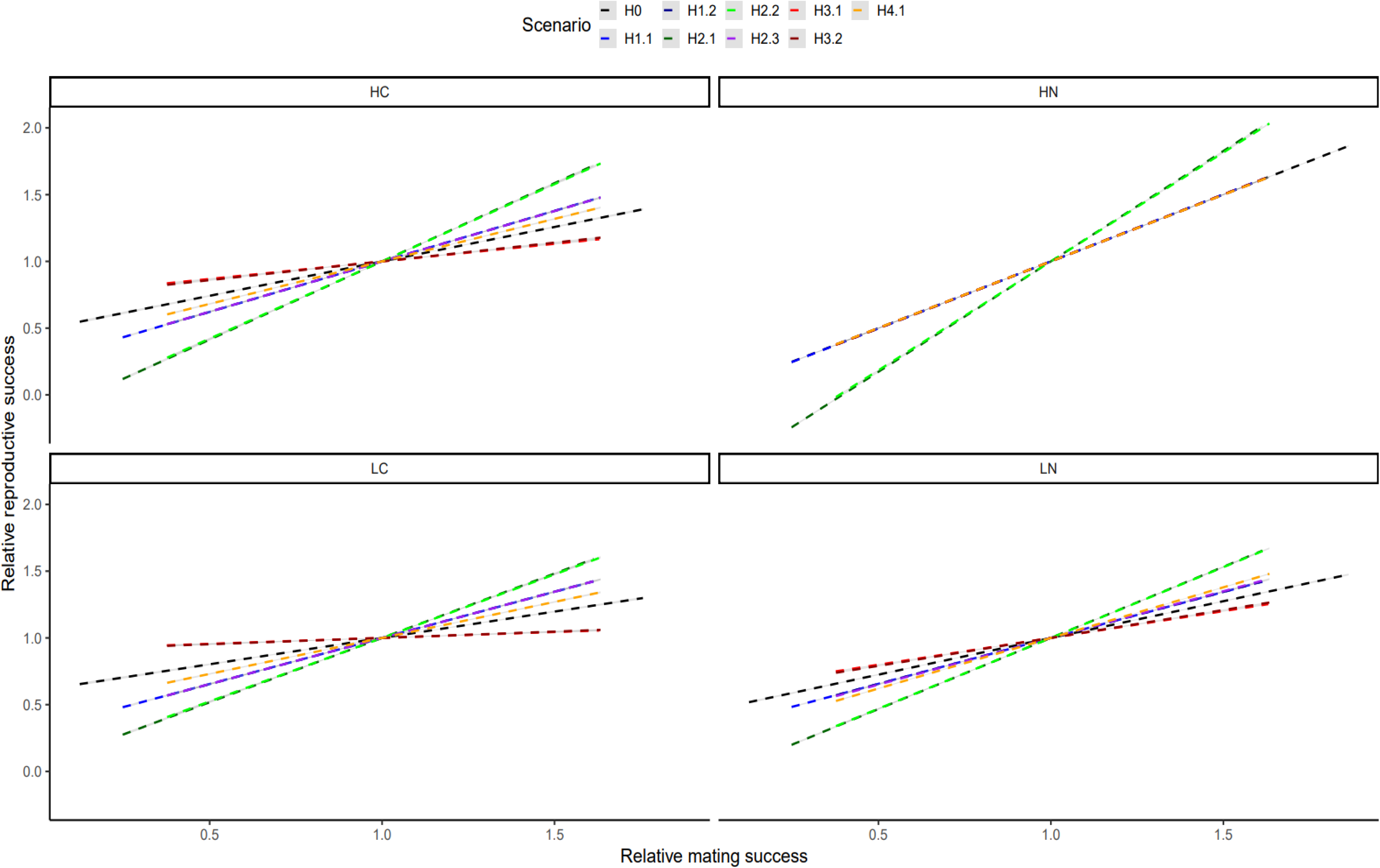
OLS linear regression between *ms* and *rs*, for each of the different scenarios modelled in each of the four systems (i.e. low or high anisogamy; presence or absence of sperm competition). Grey shaded area shows 95% C.I. H0: no co-variances; H1.1: positive co-variance between MS and ejaculate size; H1.2: positive co-variance between MS and sperm potency; H2.1: positive co-variance between MS and female egg number; H2.2: positive co-variance between MS and female egg allocation; H2.3: positive co-variance between MS and female sperm retention; H3.1: negative co-variance between MS and ejaculate size; H3.2: negative co-variance between MS and sperm potency; H4.1: negative co-variance between MS and proportion of remaining sperm allocated to female. HC: High anisogamy with sperm competition; HN: High anisogamy without sperm competition; LC: Low anisogamy with sperm competition; LN: Low anisogamy without sperm competition.

**Figure S8:**
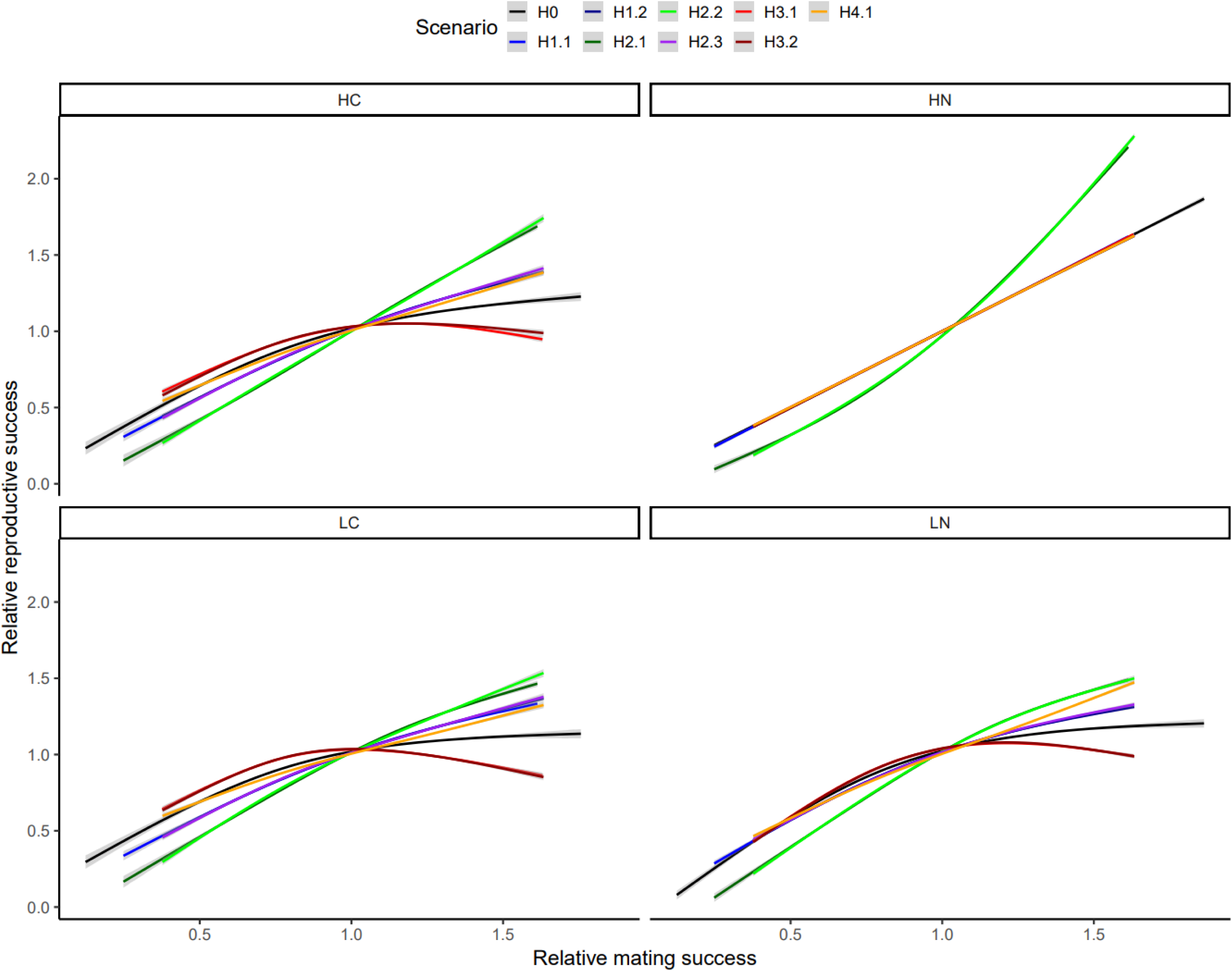
Curvilinear relationship between *ms* and *rs* for each of the nine different scenarios in each biological system, plotted as a smooth function (cubic-spline gam with four knots in ggplot). Panels show systems with different degrees of anisogamy (low or high) and different degrees of sperm competition (present or absent). H0:null shows the BG from the null scenario where there are no co-variances between MS and any other variable, therefore the influence of *ms* on *rs* is causal. H1.1 to H4.1 show BG from models where MS co-varies with other male or female traits. HC: High anisogamy with sperm competition; HN: High anisogamy without sperm competition; LC: Low anisogamy with sperm competition; LN: Low anisogamy without sperm competition.

**Figure S9:**
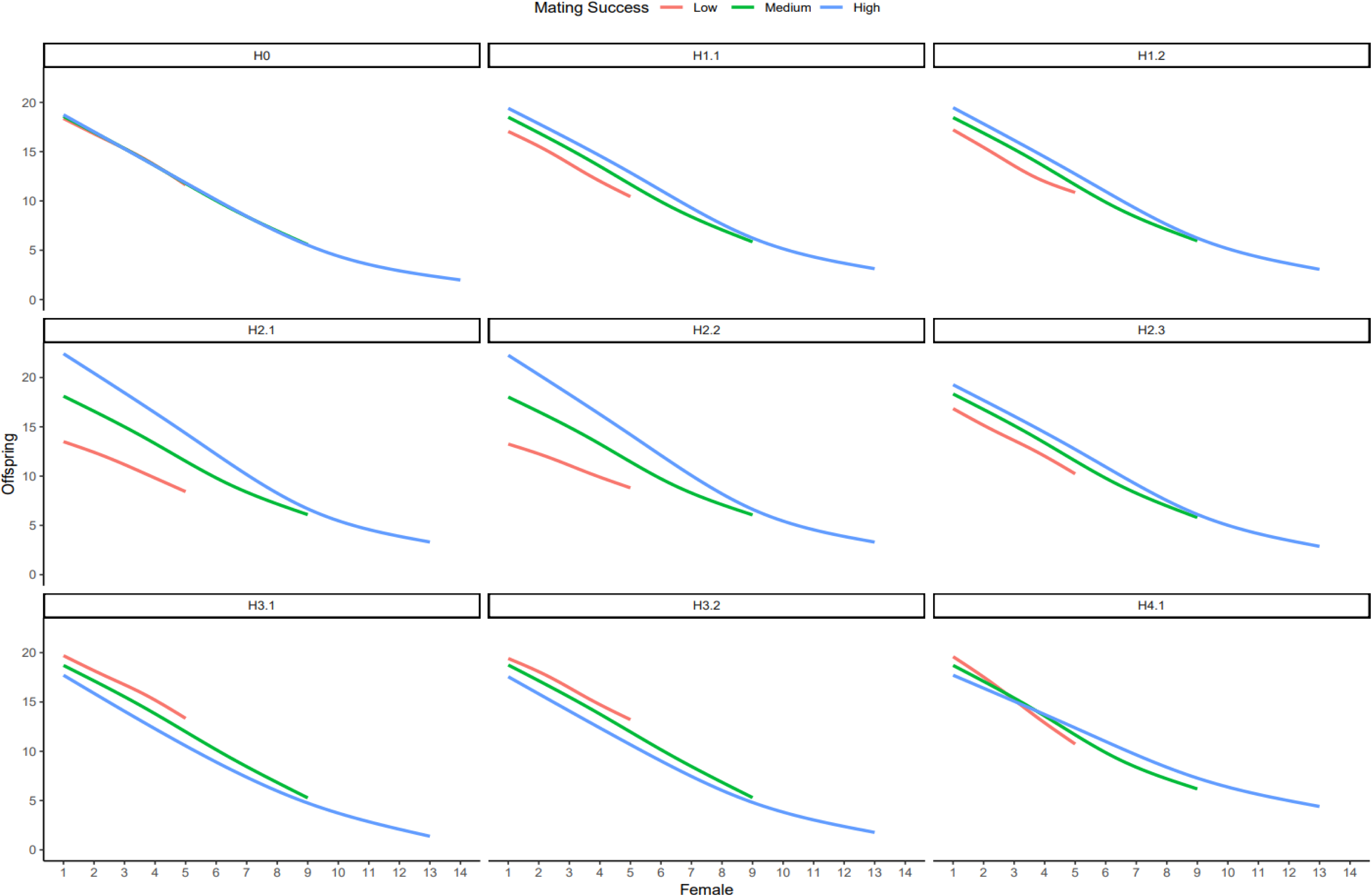
Effect of MS on the number of offspring a focal male produces with each female (rank) in his mating sequence. Panel labels correspond to each scenario in the system with high anisogamy and sperm competition. MS binned into three categories for ease of visualisation. Lines show means of 20 replicates, constructed as gam smooths with four knots in ggplot. H0: no co-variances; H1.1: positive co-variance between MS and ejaculate size (*Q*); H1.2: positive co-variance between MS and sperm potency (*V*); H2.1: positive co-variance between MS and female egg number (*F*); H2.2: positive co-variance between MS and female egg allocation (*P*); H2.3: positive co-variance between MS and female sperm retention (*L*); H3.1: negative co-variance between MS and ejaculate size (*Q*); H3.2: negative co-variance between MS and sperm potency (*V*); H4.1: negative co-variance between MS and proportion of stored ejaculate transferred by male to female (*T*).

### Supplementary tables

**Table S1:**
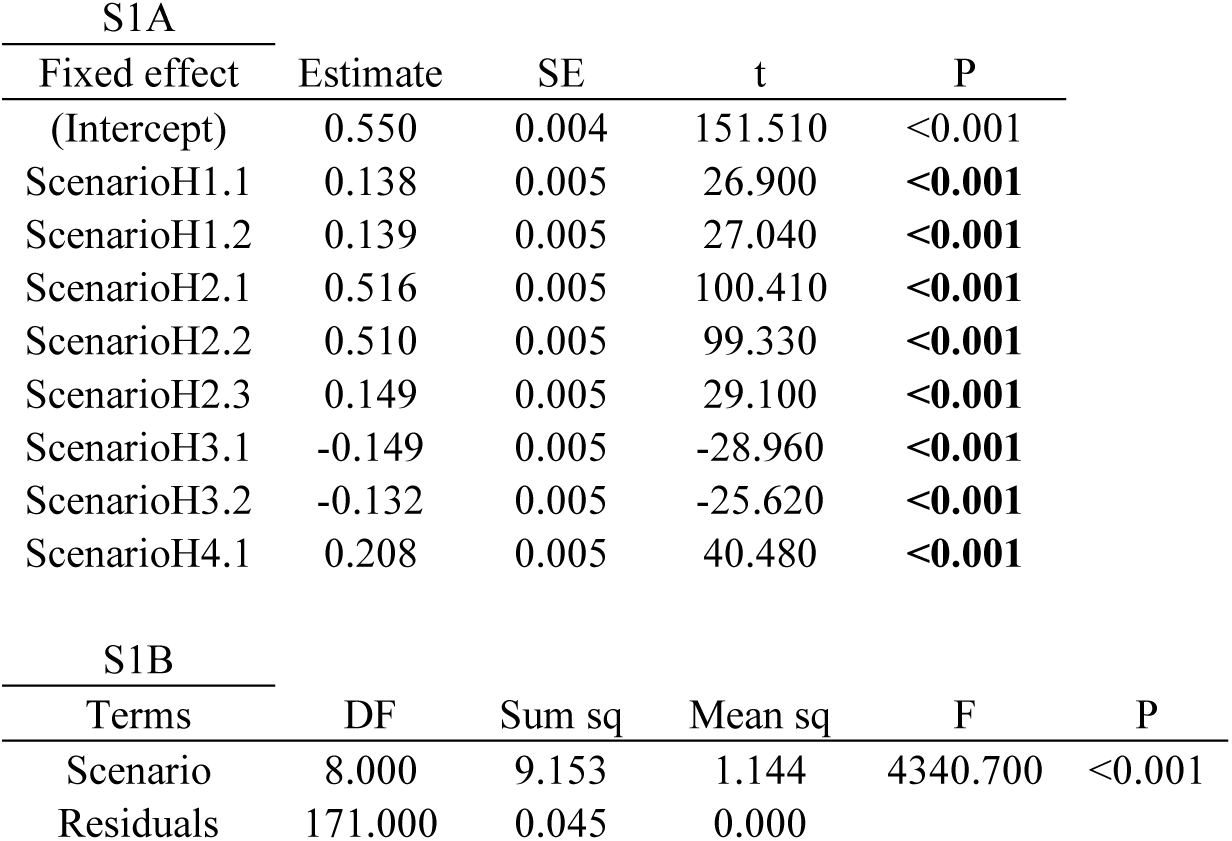
A- Linear model testing for differences in BG between the null scenario and each of the co-variance scenarios, in the low anisogamy system without sperm competition. The intercept is the BG of the null scenario (H0). The estimate is the difference in the BG between the null scenario and co-variance scenario. Sample size is 20 values (i.e. from 20 replicates) per scenario. B- ANOVA on the linear model in Table S1A, to test for overall differences in means between different scenarios. Significant differences shown in bold.

**Table S2:**
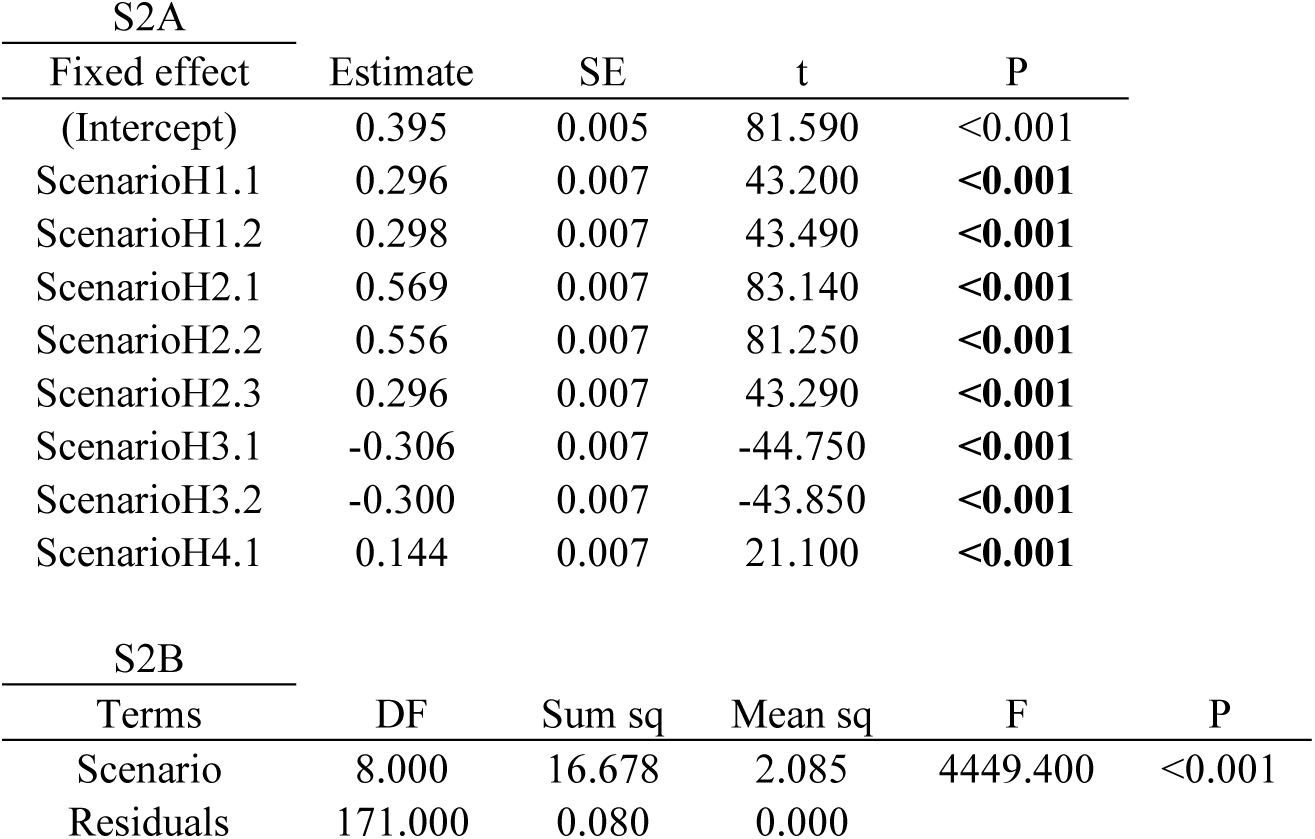
A- Linear model testing for differences in BG between the null scenario and each of the co-variance scenarios, in the low anisogamy system with sperm competition. The intercept is the BG of the null scenario (H0). The estimate is the difference in the BG between the null scenario and co-variance scenario. Sample size is 20 values (i.e. from 20 replicates) per scenario. B- ANOVA on the linear model in Table S2A, to test for overall differences in means between different scenarios. Significant differences shown in bold.

**Table S3:**
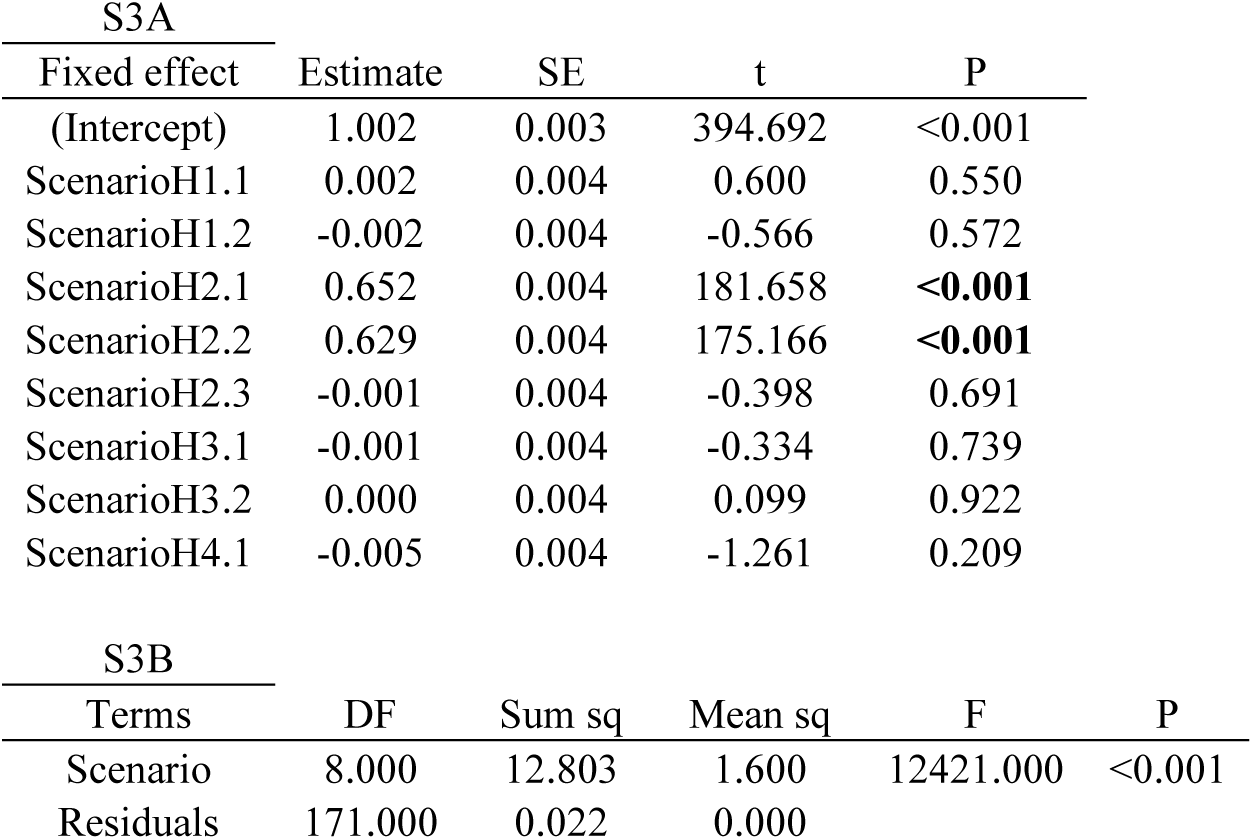
A- Linear model testing for differences in BG between the null scenario and each of the co-variance scenarios, in the high anisogamy system without sperm competition. The intercept is the BG of the null scenario (H0). The estimate is the difference in the BG between the null scenario and co-variance scenario. Sample size is 20 values (i.e. from 20 replicates) per scenario. B- ANOVA on the linear model in Table S3A, to test for overall differences in means between different scenarios. Significant differences shown in bold.

**Table S4:**
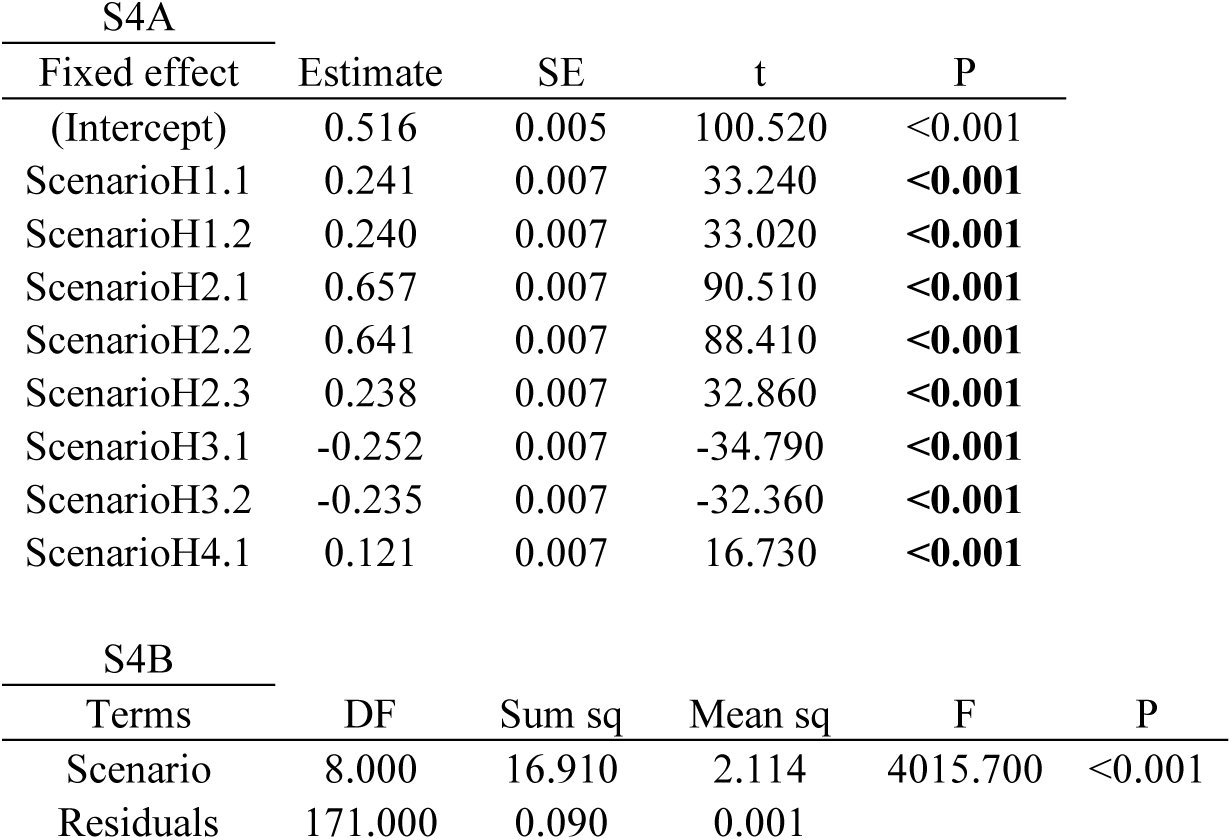
A- Linear model testing for differences in BG between the null scenario and each of the co-variance scenarios, in the high anisogamy system with sperm competition. The intercept is the BG of the null scenario (H0). The estimate is the difference in the BG between the null scenario and co-variance scenario. Sample size is 20 values (i.e. from 20 replicates) per scenario. B- ANOVA on the linear model in Table S4A, to test for overall differences in means between different scenarios. Significant differences shown in bold.

