## Supplementary information for "Simulations provide a mechanistic interpretation of Bateman gradients"

### Supporting information

#### Supplementary Figures

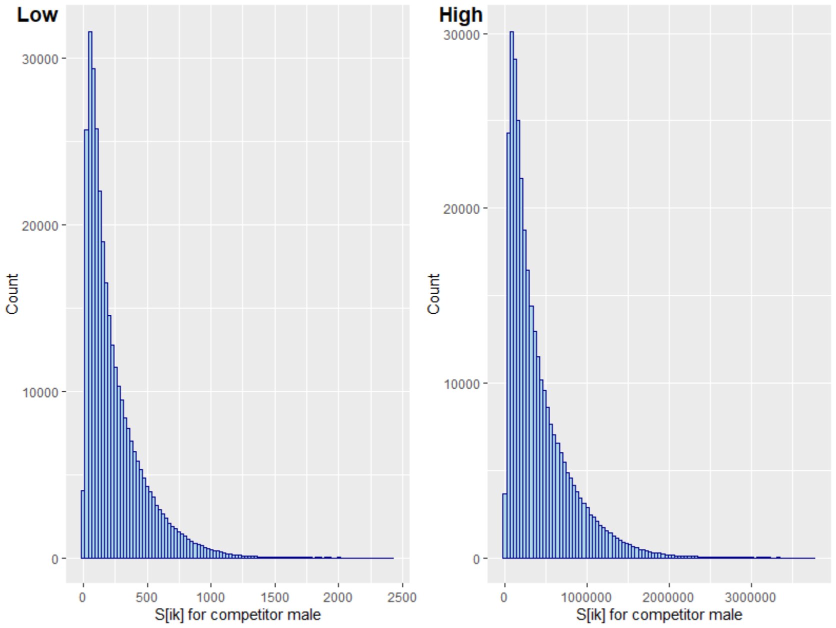


Figure S1: Distribution of $S_{ik}$ values for the competitor male in low and high anisogamy systems, from which $A_{ik}$ values for the competitor male's sperm are sampled. $S_{ik}$ represents the numbers of potent sperm transferred by a focal male and retained by a mated female, if she mated with a randomly chosen focal male at a random point in his mating sequence.

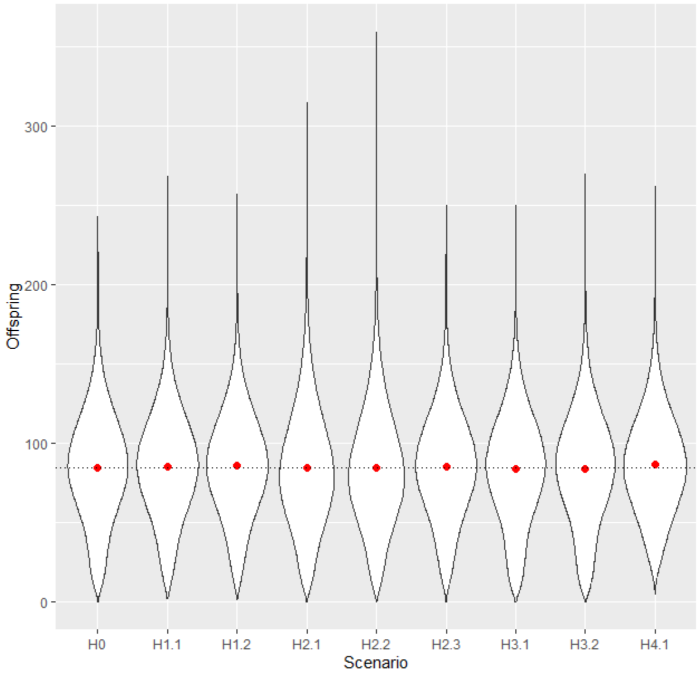


Figure S2: Distribution of offspring production of the focal male with each mated female, across the nine simulated scenarios, in the system with low anisogamy and no sperm competition. Offspring production was similar across scenarios, therefore differences in BG across scenarios cannot be attributed to difference in mean offspring production. Violin plots represent distribution of offspring produced by each focal male with each female, red dot represents mean offspring number per scenario. Dotted line shows overall mean across all scenarios. Data pooled from 20 replicates.

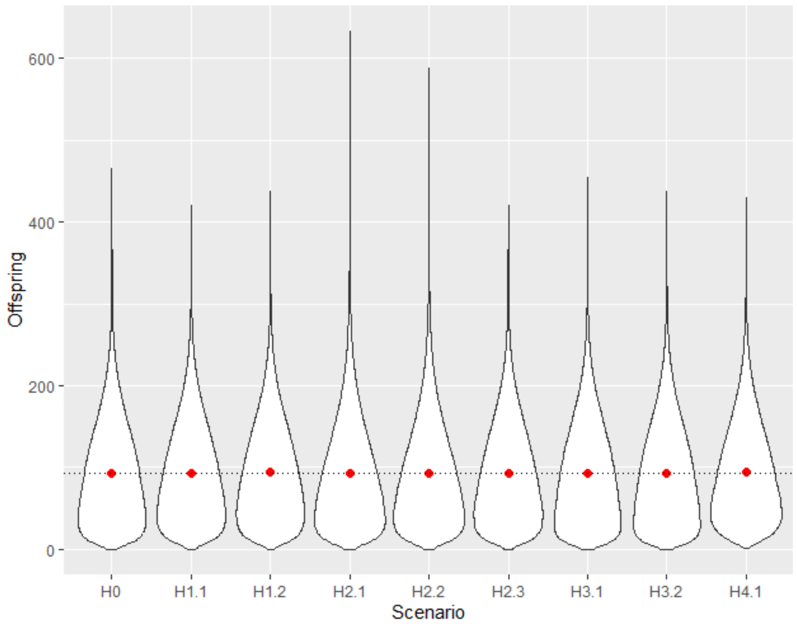


Figure S3: Distribution of offspring production of the focal male with each mated female, across the nine simulated scenarios, in the system with low anisogamy with sperm competition. Offspring production was similar across scenarios, therefore differences in BG across scenarios cannot be attributed to difference in mean offspring production. Violin plots represent distribution of offspring produced by each focal male with each female, red dot represents mean offspring number per scenario. Dotted line shows overall mean across all scenarios. Data pooled from 20 replicates.

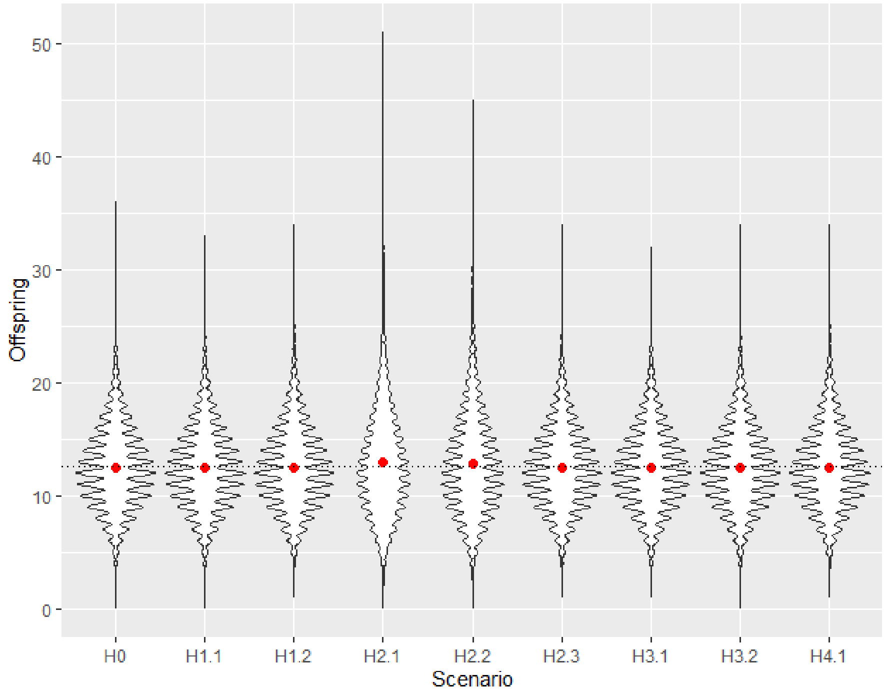


Figure S4: Distribution of offspring production of the focal male with each mated female, across the nine simulated scenarios, in the high anisogamy system without sperm competition. Offspring production was similar across scenarios, therefore differences in BG across scenarios cannot be attributed to difference in mean offspring production. Violin plots represent distribution of offspring produced by each focal male with each female, yellow triangle represents mean offspring number per scenario. Dotted line shows overall mean across all scenarios. Data pooled from 20 replicates.

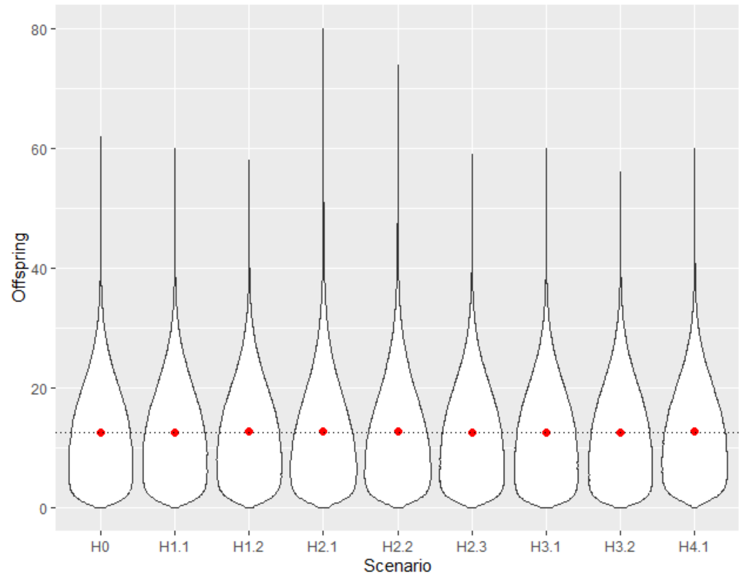


Figure S5: Distribution of offspring production of the focal male with each mated female, across the nine simulated scenarios, in the high anisogamy system with sperm competition. Offspring production was similar across scenarios, therefore differences in BG across scenarios cannot be attributed to difference in mean offspring production. Violin plots represent distribution of offspring produced by each focal male with each female, red dot represents mean offspring number per scenario. Dotted line shows overall mean across all scenarios. Data pooled from 20 replicates.

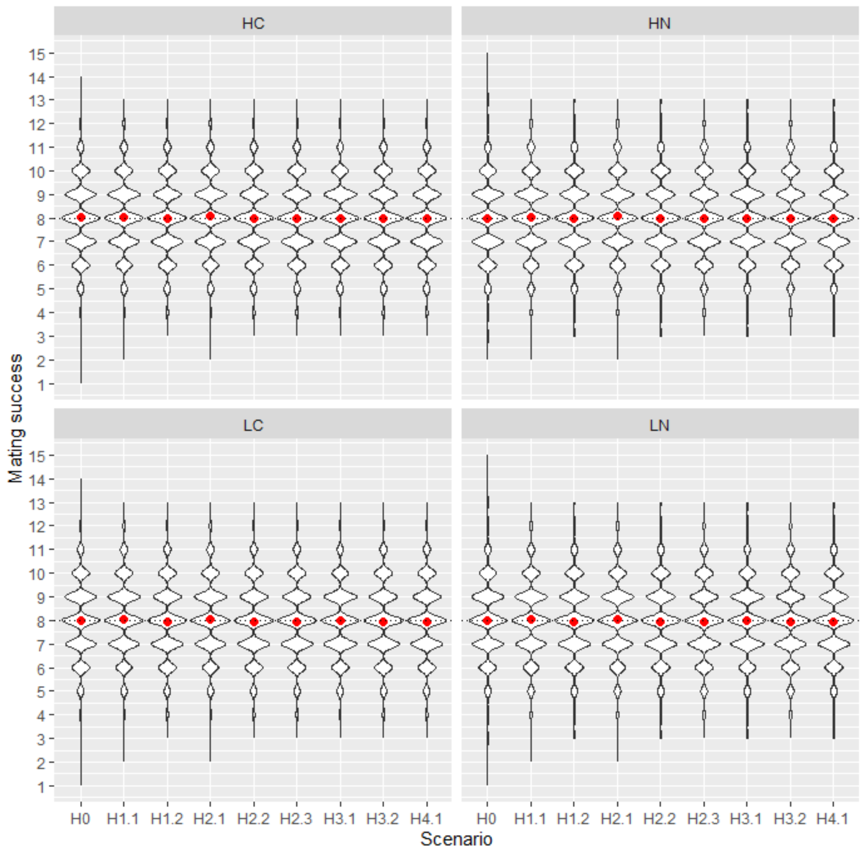


Figure S6: Distribution of male mating success across the nine scenarios and 4 systems. Note that no males in our simulations had a mating success of <1 or >15. High (H) or low anisogamy (L), with (C) or without (N) sperm competition.

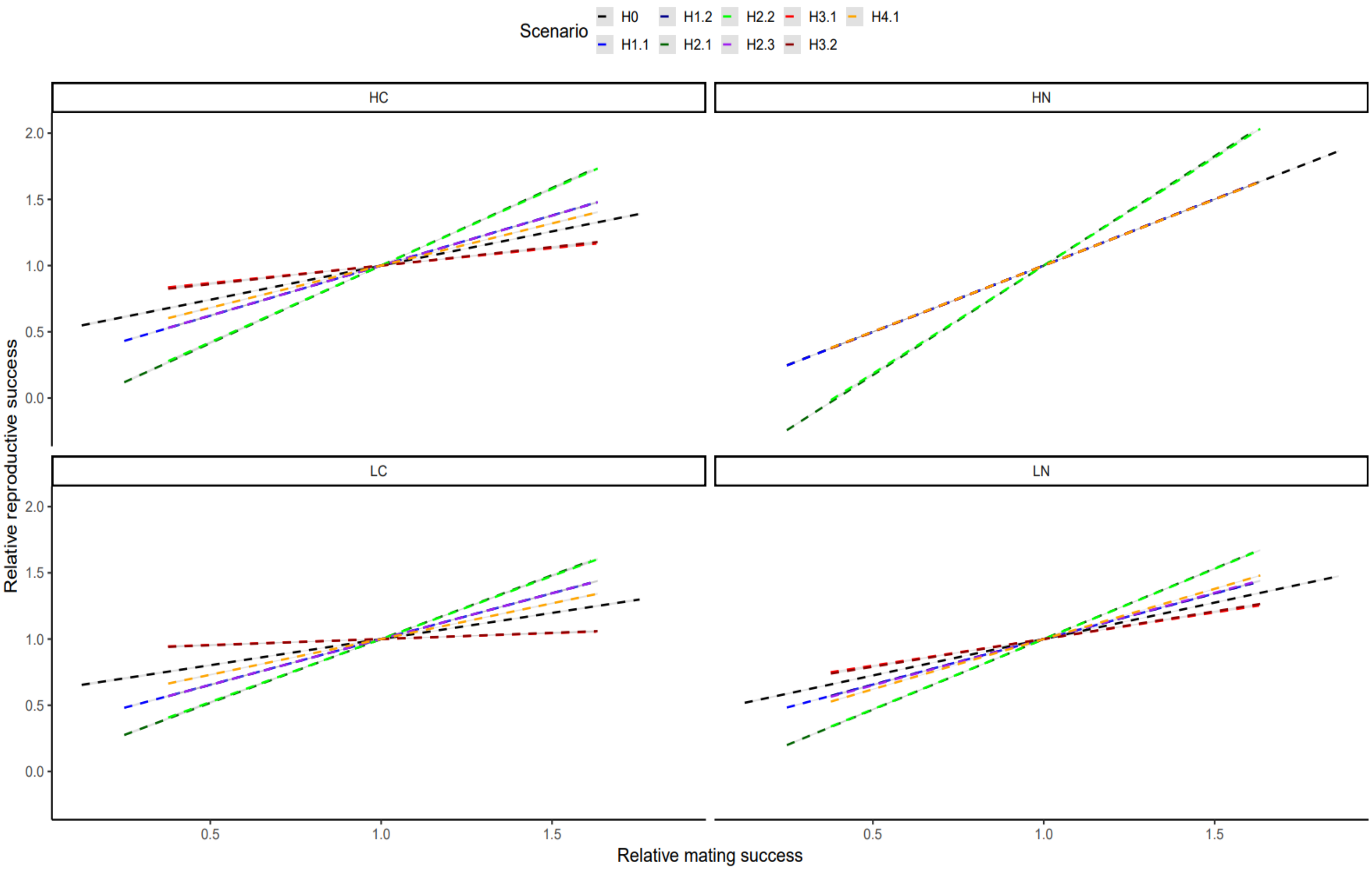
Scenario
H0
H1.2
H2.2
H3.1
H4.1
H1.1
H2.1
H2.3
H3.2
HC
HN
LC
LN
Relative reproductive success
2.0
1.5
1.0
0.5
0.0
Relative mating success
0.5
1.0
1.5

42

Figure S7: OLS linear regression between *ms* and *rs*, for each of the different scenarios modelled in each of the four systems (i.e. low or high anisogamy; presence or absence of sperm competition). Grey shaded area shows 95% C.I. H0: no co-variances; H1.1: positive co-variance between MS and ejaculate size; H1.2: positive co-variance between MS and sperm potency; H2.1: positive co-variance between MS and female egg number; H2.2: positive co-variance between MS and female egg allocation; H2.3: positive co-variance between MS and female sperm retention; H3.1: negative co-variance between MS and ejaculate size; H3.2: negative co-variance between MS and sperm potency; H4.1: negative co-variance between MS and proportion of remaining sperm allocated to female. HC: High anisogamy with sperm competition; HN: High anisogamy without sperm competition; LC: Low anisogamy with sperm competition; LN: Low anisogamy without sperm competition.

52

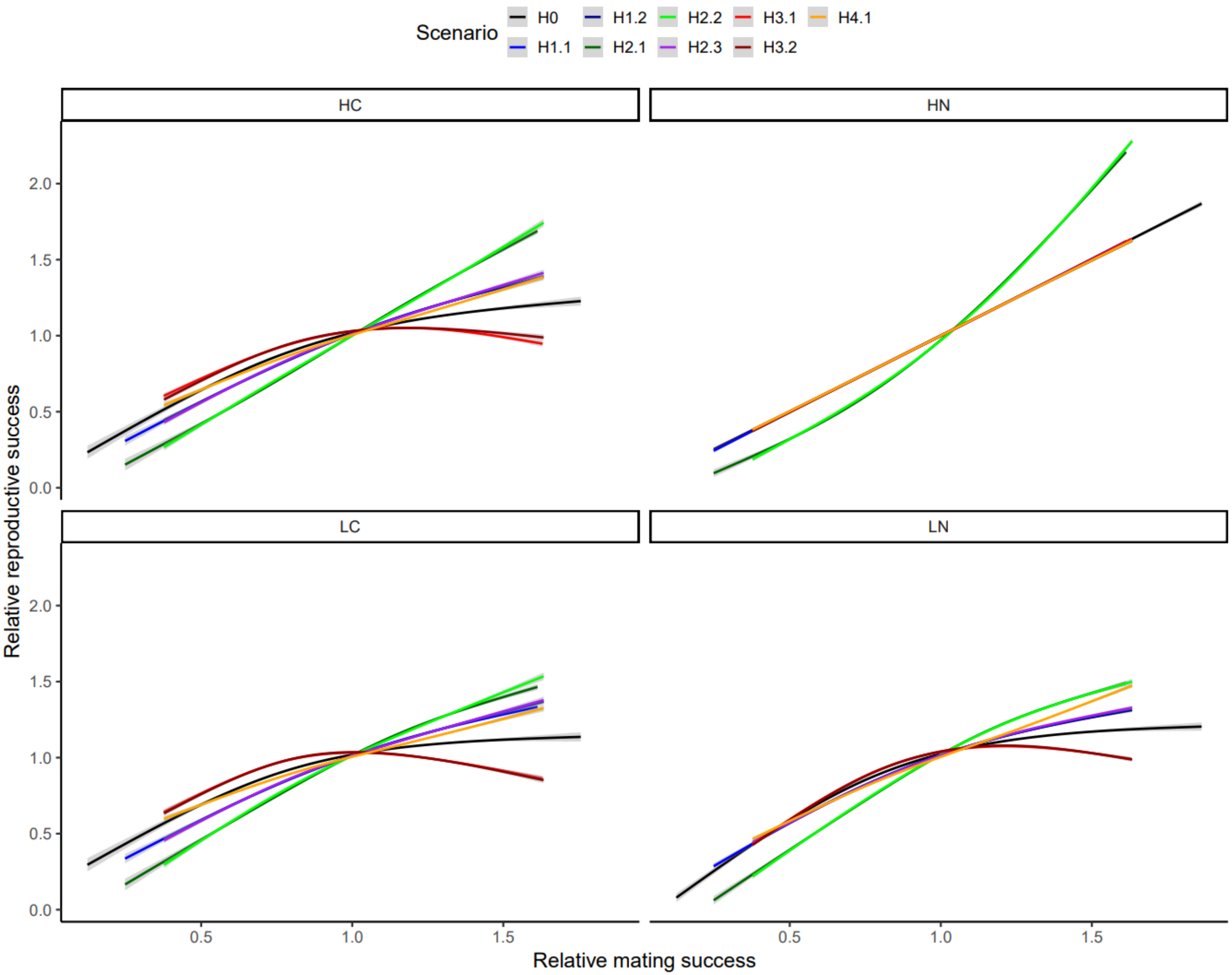

Scenario
H0
H1.1
H1.2
H2.1
H2.2
H2.3
H3.1
H3.2
H4.1
HC
HN
LC
LN
Relative reproductive success
Relative mating success
0.0
0.5
1.0
1.5
2.0

Figure S8: Curvilinear relationship between *ms* and *rs* for each of the nine different scenarios in each biological system, plotted as a smooth function (cubic-spline gam with four knots in ggplot). Panels show systems with different degrees of anisogamy (low or high) and different degrees of sperm competition (present or absent). H0:null shows the BG from the null scenario where there are no co-variances between MS and any other variable, therefore the influence of *ms* on *rs* is causal. H1.1 to H4.1 show BG from models where MS co-varies with other male or female traits. HC: High anisogamy with sperm competition; HN: High anisogamy without sperm competition; LC: Low anisogamy with sperm competition; LN: Low anisogamy without sperm competition.

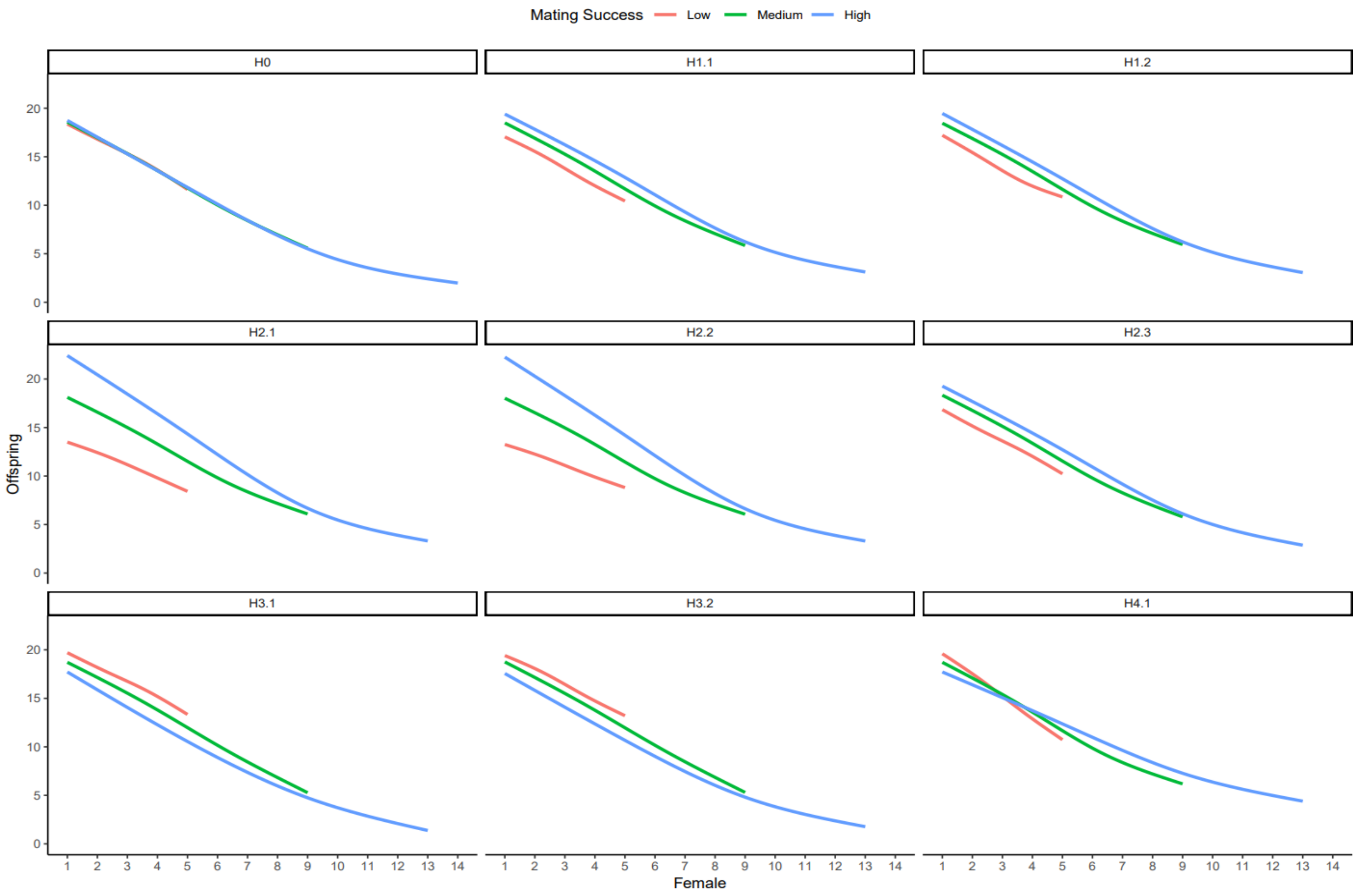
Mating Success
Low
Medium
High
H0
H1.1
H1.2
H2.1
H2.2
H2.3
H3.1
H3.2
H4.1
Offspring
0
5
10
15
20
1 2 3 4 5 6 7 8 9 10 11 12 13 14
Female

64

Figure S9: Effect of MS on the number of offspring a focal male produces with each female (rank) in his mating sequence. Panel labels correspond to each scenario in the system with high anisogamy and sperm competition. MS binned into three categories for ease of visualisation. Lines show means of 20 replicates, constructed as gam smooths with four knots in ggplot. H0: no co-variances; H1.1: positive co-variance between MS and ejaculate size (*Q*); H1.2: positive co-variance between MS and sperm potency (*V*); H2.1: positive co-variance between MS and female egg number (*F*); H2.2: positive co-variance between MS and female egg allocation (*P*); H2.3: positive co-variance between MS and female sperm retention (*L*); H3.1: negative co-variance between MS and ejaculate size (*Q*); H3.2: negative co-variance between MS and sperm potency (*V*); H4.1: negative co-variance between MS and proportion of stored ejaculate transferred by male to female (*T*).

#### Supplementary tables

Table S1: A- Linear model testing for differences in BG between the null scenario and each of the co-variance scenarios, in the low anisogamy system without sperm competition. The intercept is the BG of the null scenario (H0). The estimate is the difference in the BG between the null scenario and co-variance scenario. Sample size is 20 values (i.e. from 20 replicates) per scenario. B- ANOVA on the linear model in Table S1A, to test for overall differences in means between different scenarios. Significant differences shown in bold.

S1A

| Fixed effect | Estimate | SE | t | P |
|---|---|---|---|---|
| (Intercept) | 0.550 | 0.004 | 151.510 | <0.001 |
| ScenarioH1.1 | 0.138 | 0.005 | 26.900 | **<0.001** |
| ScenarioH1.2 | 0.139 | 0.005 | 27.040 | **<0.001** |
| ScenarioH2.1 | 0.516 | 0.005 | 100.410 | **<0.001** |
| ScenarioH2.2 | 0.510 | 0.005 | 99.330 | **<0.001** |
| ScenarioH2.3 | 0.149 | 0.005 | 29.100 | **<0.001** |
| ScenarioH3.1 | -0.149 | 0.005 | -28.960 | **<0.001** |
| ScenarioH3.2 | -0.132 | 0.005 | -25.620 | **<0.001** |
| ScenarioH4.1 | 0.208 | 0.005 | 40.480 | **<0.001** |

S1B

| Terms | DF | Sum sq | Mean sq | F | P |
|---|---|---|---|---|---|
| Scenario | 8.000 | 9.153 | 1.144 | 4340.700 | <0.001 |
| Residuals | 171.000 | 0.045 | 0.000 | | |

Table S2: A- Linear model testing for differences in BG between the null scenario and each of the co-variance scenarios, in the low anisogamy system with sperm competition. The intercept is the BG of the null scenario (H0). The estimate is the difference in the BG between the null scenario and co-variance scenario. Sample size is 20 values (i.e. from 20 replicates) per scenario. B- ANOVA on the linear model in Table S2A, to test for overall differences in means between different scenarios. Significant differences shown in bold.

S2A

| Fixed effect | Estimate | SE | t | P |
|---|---|---|---|---|
| (Intercept) | 0.395 | 0.005 | 81.590 | <0.001 |
| ScenarioH1.1 | 0.296 | 0.007 | 43.200 | **<0.001** |
| ScenarioH1.2 | 0.298 | 0.007 | 43.490 | **<0.001** |
| ScenarioH2.1 | 0.569 | 0.007 | 83.140 | **<0.001** |
| ScenarioH2.2 | 0.556 | 0.007 | 81.250 | **<0.001** |
| ScenarioH2.3 | 0.296 | 0.007 | 43.290 | **<0.001** |
| ScenarioH3.1 | -0.306 | 0.007 | -44.750 | **<0.001** |
| ScenarioH3.2 | -0.300 | 0.007 | -43.850 | **<0.001** |
| ScenarioH4.1 | 0.144 | 0.007 | 21.100 | **<0.001** |

S2B

| Terms | DF | Sum sq | Mean sq | F | P |
|---|---|---|---|---|---|
| Scenario | 8.000 | 16.678 | 2.085 | 4449.400 | <0.001 |
| Residuals | 171.000 | 0.080 | 0.000 | | |

Table S3: A- Linear model testing for differences in BG between the null scenario and each of the co-variance scenarios, in the high anisogamy system without sperm competition. The intercept is the BG of the null scenario (H0). The estimate is the difference in the BG between the null scenario and co-variance scenario. Sample size is 20 values (i.e. from 20 replicates) per scenario. B- ANOVA on the linear model in Table S3A, to test for overall differences in means between different scenarios. Significant differences shown in bold.

| S3A | | | | |
|---|---|---|---|---|
| Fixed effect | Estimate | SE | t | P |
| (Intercept) | 1.002 | 0.003 | 394.692 | <0.001 |
| ScenarioH1.1 | 0.002 | 0.004 | 0.600 | 0.550 |
| ScenarioH1.2 | -0.002 | 0.004 | -0.566 | 0.572 |
| ScenarioH2.1 | 0.652 | 0.004 | 181.658 | **<0.001** |
| ScenarioH2.2 | 0.629 | 0.004 | 175.166 | **<0.001** |
| ScenarioH2.3 | -0.001 | 0.004 | -0.398 | 0.691 |
| ScenarioH3.1 | -0.001 | 0.004 | -0.334 | 0.739 |
| ScenarioH3.2 | 0.000 | 0.004 | 0.099 | 0.922 |
| ScenarioH4.1 | -0.005 | 0.004 | -1.261 | 0.209 |

| S3B | | | | | |
|---|---|---|---|---|---|
| Terms | DF | Sum sq | Mean sq | F | P |
| Scenario | 8.000 | 12.803 | 1.600 | 12421.000 | <0.001 |
| Residuals | 171.000 | 0.022 | 0.000 | | |

Table S4: A- Linear model testing for differences in BG between the null scenario and each of the co-variance scenarios, in the high anisogamy system with sperm competition. The intercept is the BG of the null scenario (H0). The estimate is the difference in the BG between the null scenario and co-variance scenario. Sample size is 20 values (i.e. from 20 replicates) per scenario. B- ANOVA on the linear model in Table S4A, to test for overall differences in means between different scenarios. Significant differences shown in bold.

| S4A | | | | |
|---|---|---|---|---|
| Fixed effect | Estimate | SE | t | P |
| (Intercept) | 0.516 | 0.005 | 100.520 | <0.001 |
| ScenarioH1.1 | 0.241 | 0.007 | 33.240 | **<0.001** |
| ScenarioH1.2 | 0.240 | 0.007 | 33.020 | **<0.001** |
| ScenarioH2.1 | 0.657 | 0.007 | 90.510 | **<0.001** |
| ScenarioH2.2 | 0.641 | 0.007 | 88.410 | **<0.001** |
| ScenarioH2.3 | 0.238 | 0.007 | 32.860 | **<0.001** |
| ScenarioH3.1 | -0.252 | 0.007 | -34.790 | **<0.001** |
| ScenarioH3.2 | -0.235 | 0.007 | -32.360 | **<0.001** |
| ScenarioH4.1 | 0.121 | 0.007 | 16.730 | **<0.001** |

| S4B | | | | | |
|---|---|---|---|---|---|
| Terms | DF | Sum sq | Mean sq | F | P |
| Scenario | 8.000 | 16.910 | 2.114 | 4015.700 | <0.001 |
| Residuals | 171.000 | 0.090 | 0.001 | | |
